# Physical and transcriptional organisation of the bread wheat intracellular immune receptor repertoire

**DOI:** 10.1101/339424

**Authors:** Burkhard Steuernagel, Kamil Witek, Simon G. Krattinger, Ricardo H. Ramirez-Gonzalez, Henk-jan Schoonbeek, Guotai Yu, Erin Baggs, Agnieszka I. Witek, Inderjit Yadav, Ksenia V. Krasileva, Jonathan D. G. Jones, Cristobal Uauy, Beat Keller, Christopher J. Ridout, The International Wheat Genome Sequencing Consortium, Brande B. H. Wulff

## Abstract

Disease resistance genes encoding intracellular immune receptors of the nucleotide-binding and leucine-rich repeat (NLR) class of proteins detect pathogens by the presence of pathogen effectors. Plant genomes typically contain hundreds of NLR encoding genes. The availability of the hexaploid wheat cultivar Chinese Spring reference genome now allows a detailed study of its NLR complement. However, low NLR expression as well as high intra-family sequence homology hinders their accurate gene annotation. Here we developed NLR-Annotator for *in silico* NLR identification independent of transcript support. Although developed for wheat, we demonstrate the universal applicability of NLR-Annotator across diverse plant taxa. Applying our tool to wheat and combining it with a transcript-validated subset of genes from the reference gene annotation, we characterized the structure, phylogeny and expression profile of the NLR gene family. We detected 3,400 full-length NLR loci of which 1,540 were confirmed as complete genes. NLRs with integrated domains mostly group in specific sub-clades. Members of another subclade predominantly locate in close physical proximity to NLRs carrying integrated domains suggesting a paired helper-function. Most NLRs (88%) display low basal expression (in the lower 10 percentile of transcripts), which may be tissue-specific and/or induced by biotic stress. As a case study for applying our tool to the positional cloning of resistance genes, we estimated the number of NLR genes within the intervals of mapped rust resistance genes. Our study will support the identification of functional resistance genes in wheat to accelerate the breeding and engineering of disease resistant varieties.

## Background

The status of wheat as the world’s most widely grown and important food crop [1] is threatened by the emergence and spread of new and old diseases. For example, wheat stem rust, long considered a vanquished foe of the past, has in the last 20 years caused devastating epidemics in Africa [2, 3] and eastern Russia [4], while, in 2013 and 2016 large outbreaks occurred for the first time in >50 years in western Europe [5, 6]. The spread of disease into new regions can be attributed to the very success of wheat as a globally traded commodity. In that context, wheat blast, a new disease for wheat, which had until recently been confined to Brazil and other countries in South America, appearead in 2016 in Bangladesh, possibly as a result of importing contaminated grain [7, 8]. The warm, wet climates in the wheat belts of India and China, which supply ∼30% of the worlds wheat [1], favour the further proliferation of this devastating disease. In temperate regions, Septoria tritici blotch has become a major foliar disease of wheat, which with the emergence of fungicide resistant strains [9, 10] and the forthcoming restrictions and bans on the remaining effective chemicals [11], threatens to cause almost complete collapse of wheat production in some regions [12]. The breeding of new wheat cultivars with improved genetic resistance to a diverse array of pathogens is therefore of great importance to ensure future wheat production.

In plants, heritable genetic variation for disease resistance is often controlled by dominant resistance (R) genes encoding intracellular immune receptors with nucleotide-binding and leucine-rich repeat (NLR) domains [13]. NLRs detect the presence of pathogen effector molecules which are delivered into the plant cell by the pathogen to promote virulence [14]. NLRs either detect effectors directly or more frequently, indirectly through the modification of a host target by the pathogen effector. In the latter case, the NLR is often said to be ‘guarding’ the pathogenicity target [15]. Some NLRs contain an integrated domain, which has been proposed to act as a decoy to the intended effector pathogenicity target [16–18].

NLRs belong to one of the largest multi-gene families in plants. A plant genome may contain several hundred NLRs [16]. Many NLRs are under extreme diversifying selection, to the extent that two accessions from the same species can display significant NLR copy number and sequence variation, the result of the selection of new variants based on duplication, deletion, unequal crossing over and mutation [19–22]. In wheat, 15 *R* genes for resistance to wheat rusts and powdery mildew have been cloned that encode NLRs [23–34].

A major impediment to studying NLRs in wheat and the cloning of functional *R* genes, has been the lack of a contiguous reference genome sequence. Although the genome of bread wheat (*Triticum aestivum*) is one of the most challenging crop genomes to study due to its large size (15.4-15.8 Gb) [35], repetitiveness (∼85%), and hexaploid nature, recent technological advances [36] as well as a strong interest by governmental and industrial funding sources have allowed completion of the genome sequence. Starting in 2012 with a low coverage survey sequence [37], improved assemblies underpinned by different strategies have been published [38–41]. In the beginning of 2017, the first version of a wheat genome sequence with chromosome-sized scaffolds (IWGSC RefSeq v1.0), was made publicly available. The major results of this project are summarized in the study by the International Wheat Genome Sequencing Consortium [35], and have resulted in a large number of additional studies including the analysis of the transcriptional landscape of wheat [42]. In addition, high quality reference genome sequences have been recently published for *Aegilops tauschii* [43, 44], the wild diploid progenitor of the wheat D genome, and wild emmer (*T. diccocoides*) [45], the wild AB tetraploid progenitor.

The new high quality wheat and wild wheat reference genomes will undoubtedly facilitate the study of wheat NLR structure, function and evolution. However, the study of NLRs, including those in wheat, is complicated by (i) the absence of a tool for accurate and high-throughput annotation, and (ii) the extremely low basal level of expression for many NLR genes. To overcome these two additional challenges, we firstly developed NLR-Annotator, a tool for *de novo* genome annotation of loci associated with NLRs. Secondly, we performed NLR exome capture and sequencing on cDNA [46] isolated from different tissues and developmental stages to obtain the expression profile and intron/exon structure of wheat NLRs. By applying NLR-Annotator and our expression data to the genome of the wheat reference cultivar Chinese Spring, we found 3,400 loci that may be functional NLR genes or pseudo genes. In a series of studies we further demonstrate the application of our tool and expression data, to show that wheat NLRs predominantly occur towards the telomeres and in close proximity to each other, display pronounced copy number and sequence variation between the A, B and D sub-genomes (yet maintain conservation of intron-exon structure), display clade-specific integration of novel domains, and that NLR expression is tissue specific and modulated by development and biotic stress.

## Results

### Motif-based annotation of NLR loci in whole genome assemblies

We set out to develop a method for annotation of NLRs independent from gene calling. The recently published pipeline NLR-Parser [47] uses combinations of short motifs of 15 to 50 amino acids to classify a sequence as NLR-related. These motifs had been defined based on manual curation of a training set of known NLR sequences by Jupe et al. [48] and mainly resemble the sub-structures of NLR protein domains also described in other studies (Figure S1; Jupe et al [48]). Since these motifs may occur randomly in a genome, the NLR-Parser searches for combinations of tuples or triplets of motifs that often occur in the same order. The drawback of NLR-Parser, however, is that it can only classify a sequence, not distinguish the border between two NLRs within the same sequence. In the extreme case of a whole chromosome with multiple NLRs, this would be classified as a single complete NLR.

Here, we present NLR-Annotator as an extension of NLR-Parser and a tool to annotate NLR loci in genomic sequence data. In this study, we define the term ‘NLR locus’ as a section of genomic sequence associated with a single NLR, i.e. one NB-ARC domain potentially followed by one or more LRRs. Our pipeline disects genomic sequences into overlapping fragments, and then uses NLR-Parser to pre-select those fragments potentially harbouring NLR loci. In this step, the nucleotide sequence of each fragment is translated in all six frames to search for motifs. Subsequently, the positions of each motif are transferred to nucleotide positions. The NLR-Annotator then integrates data from all fragments, evaluates motif positions and combinations and suggests a list of loci likely to be associated with NLRs. For an overview, see Figure S2. The general concept is to search for a triplet of consecutive motifs that are associated with the NB-ARC domain (Figure S1). This is used as a seed, which is then elongated into the pre-NB region and the LRRs by searching for additional motifs associated with those regions.

We tested the NLR-Annotator on the *Arabidopsis thaliana* genome sequence using previously annotated NLR genes as a gold standard. In the Col-0 reference genome TAIR10 (http://www.arabidopsis.org), 115 genes were annotated as NLRs. Using NLR-Annotator, we found 171 loci in the TAIR10 genome assembly. Of the 115 NLR annotated genes, only eight were not overlapping with one of our loci. We manually investigated those eight protein sequences and found that only three had an NB-ARC and LRR domain. Two were ADR1 and an ADR1-like (AT1G33560.1 and AT5G66900.1, respectively) and the third was a TIR-NLR (AT4G19530.1). In those three cases, for at least one motif within the NB-ARC domain the similarity to the consensus motif sequence was below the default threshold. The ADR1 genes have been reported to be missed by NLR-Parser before [47]. The third gene was detected by NLR-Parser due to its LRR motifs only, whereas motifs in the NB-ARC domain were below the threshold. For NLR-Annotator, we decided not to report LRRs without good evidence for an NB-ARC domain due to LRRs being attached to different gene families other than NLRs. This trade-off results in a slight reduction of sensitivity to avoid false positives. Of the remaining 56 loci annotated by NLR-Annotator, all but five overlapped with genes annotated in TAIR but not yet classified as NLRs. Of these five loci, three were annotated as partial loci with a nonsense mutation within a motif, thus highly likely pseudogenes. The two remaining loci had a motif composition indicative of a complete TIR-NLR.

To further test the functionality of NLR-annotator, we annotated the published genomes of eight other plant species for NLR loci (Table S1). We found 932 loci in robusta coffee (*Coffea canephora* [49], http://coffee-genome.org), 154 loci in maize (Zea *mays* [50], http://ensembl.gramene.org), 50 in papaya *(Carica papaya* [51], https://phytozome.jgi.doe.gov), 71 in cucumber (*Cucumis sativus*, https://phytozome.jgi.doe.gov), 527 in soybean (*Glycine max* [52], https://phytozome.jgi.doe.gov), 694 in potato (*Solanum tuberosum* [53], https://solgenomics.net, PGSC_DM_v3), 284 in tomato (*Solanum lycopersicum* [54], https://solgenomics.net, SL3.00) and 342 in purple false brome (*Brachypodium distachyon* [55], https://phytozome.jgi.doe.gov). In a previously published study of tomato, 355 NLRs were identified [46]. The lower number of NLRs identified by NLR-Annotator (284) is mostly due to the higher stringency of NLR-Annotator for minimal number and organisation of motifs and a requirement for the presence of an NB-ARC domain.

Using NLR-Annotator, we screened various assemblies of hexaploid bread wheat (*Triticum aestivum*) cultivar Chinese Spring. With the annotated loci we distinguished between partial and complete loci where a complete locus contains the P-loop/motif1 that marks the start of an NB-ARC domain as well as at least one LRR associated motif (Figure S1). In the assemblies from 2014 [38] and 2015 [39], we found 12,441 and 15,315 loci respectively but only 1,050 and 1,249 were complete. The assemblies made available in 2017 by Clavijo and colleagues [40], Zimin and colleagues [41] and the IWGSC [35] have a considerably smaller number of total NLR loci (3,251, 3,568 and 3,400, respectively) but concomitantly larger number of complete NLR loci (2465, 2,731 and 2,580, respectively). Possible reasons for differences in the number of detected NLR loci include differences in the genome completeness and continuity, and assembly mistakes such as collapsed regions or falsely duplicated regions. For the analyses described in this study, we used the IWGSC reference sequence to take advantage of the pseudo-chromosomes and the gene annotation [35].

### Physical position of NLR loci

With the availability of pseudo-chromosomes, which represent 94% of the predicted wheat genome size, most NLR loci can now be placed in the physical context of their location on the chromosome. The foremost practical implication of this is to take advantage of available mapping data and other resources to accelerate the identification of functional disease resistance genes. Annotated NLR loci were found to preferentially locate at the telomeres of each chromosome (Figure 1) and cluster together. More than 400 of the 3,400 observed loci are in a proximity of less than 5 kb to another locus. Half of all loci are in a distance of less than 50 kb to another locus (Figure 1). The NLR loci are distributed over all chromosomes and the number of NLR loci per chromosome ranges between 29 (chromosome 4D) and 280 (chromosome 4A). No clear preference of NLR numbers towards one homeologue within each group could be observed (Table S2).

**Figure 1.**
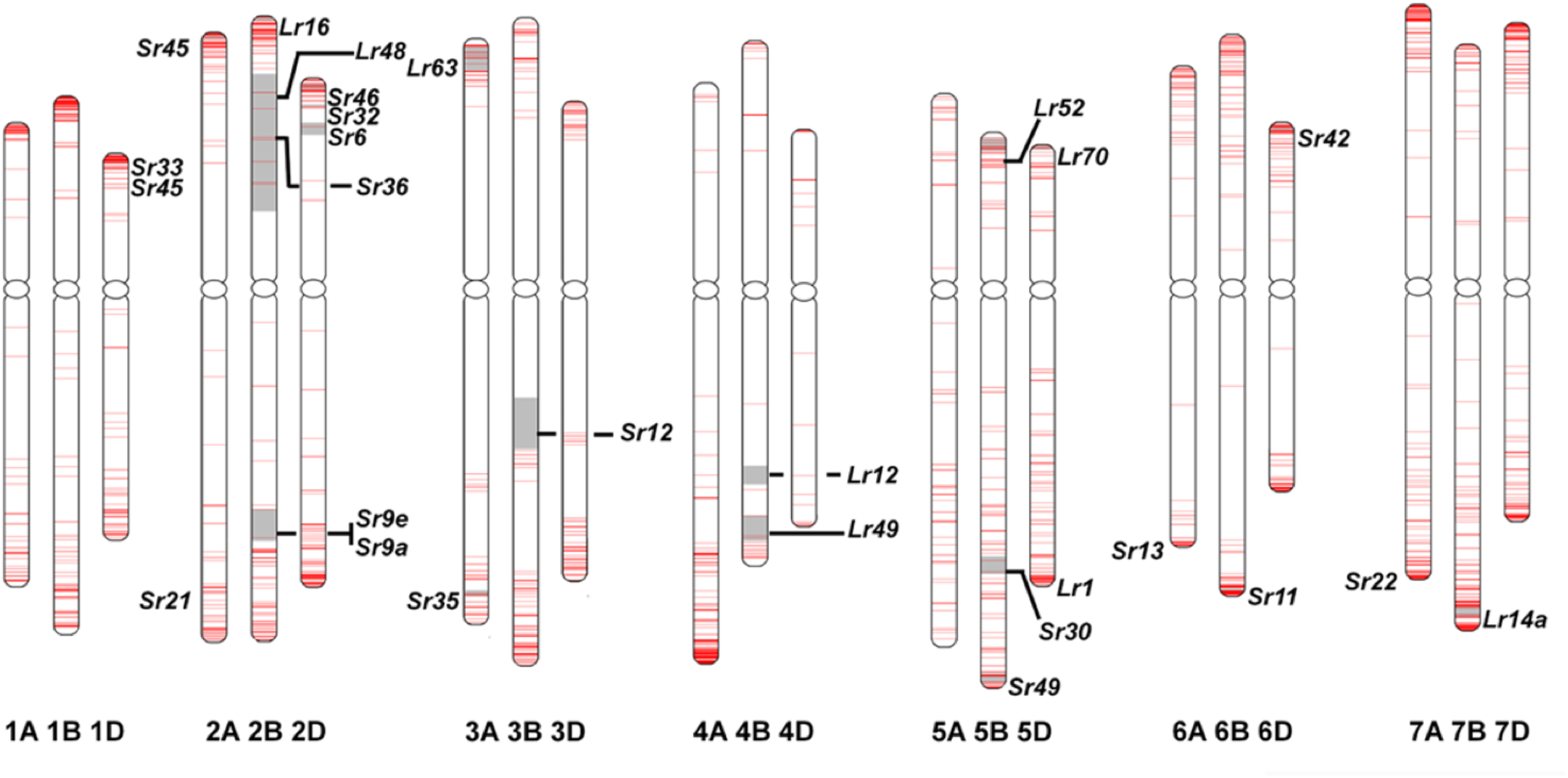
Physical position of NLR-associated loci within the wheat genome. Red bars on chromosomes mark the presence of an NLR-associated locus. Gray regions depict intervals between flanking markers of fine-mapped rust *R* genes.

To demonstrate the potential practical advantage of our resource, i.e. annotated NLR loci with a physical position on pseudo-chromosomes, we searched the literature for leaf rust and stem rust resistance genes that have been genetically mapped in wheat but not yet cloned. Most genes have been mapped in accessions other than the reference accession Chinese Spring and most resistance genes may not have a functional allele in Chinese Spring. Nevertheless, we hypothesized that in many cases, a non-functional allele or close homologue may be present, the sequence of which could then be used to speed up the cloning of the functional allele from the resistant accession.

To explore the suitability of this approach, we searched the Chinese Spring genome for homologues of several cloned *R* genes, namely *Sr22, Sr33, Sr35, Sr45, Sr50, Pm2, Pm3, Pm8, Lr1, Lr10, Lr22a, Yr5, Yr7 and YrSP* [23, 26, 27, 29–34, 56, 57]. In eight cases (*Sr22, Sr33, Sr45, Pm2, Pm3, Lr1, Yr5, Yr7*), the best alignment of the protein sequence to Chinese Spring was on the same chromosome where the gene was cloned from. For *Sr35* and *YrSp*, the second best alignment was on the right chromosome and the best alignment was on a homelologous chromosome. For *Pm8*, which was introgressed from rye, and *Lr22a*, the second best alignment was on the right chromosome whereas the best alignment on a scaffold unassigned to a chromosome yet. *Sr50* was cloned from rye as well [33] but characterized to be an orthologue of *Mla*, on group 1 chromosomes. As expected, the best three alignments for *Sr50* were therefore with chromosomes 1A, 1D and 1B. For *Lr10*, we did not find an alignment on 1A, but on the homeologous chromosomes. However, this was expected since Chinese Spring was previously characterized to be a deletion haplotype of *Lr10* [58]. Next, we aligned sequences of flanking markers from 39 mapped stem rust (Sr) and leaf rust (Lr) genes to Chinese Spring, defined the physical interval and counted the number of NLR-associated loci within the interval. The loci can be used for candidate gene approaches or further delimiting of map intervals in positional cloning projects. In 32 cases, we could determine a physical interval, either by using the exact flanking markers or by using chromosome ends or the centromere as surrogate, to determine a number of candidate NLRs. We found between 0 and 61 NLRs as candidates (average: 10) (Table S3). Interesting examples include *Sr6* (physical interval 15.8 Mb, one candidate), *Lr13* (physical interval 60 Mb, three candidates) and *Lr49* (physical interval 33 Mb, three candidates). Notably, these are large intervals with only a few candidate NLR genes, providing the opportunity to explore candidates before embarking on the more laborious tasks to further delimit or sequence the interval in the donor accession.

### Comparison of NLR loci identified by NLR-Annotator with automated gene annotation in IWGSC RefSeq v1.0

We expected a substantial number of our independently predicted NLR loci to be overlapping with genes from the automated gene annotation v1.0 of the Chinese Spring reference sequence (IWGSC RefSeq v1.0). Of the 3,400 loci we predicted by NLR-Annotator, 2,914 overlap with genes annotated in RefSeq v1.0 (2,329 of which are high-confidence (HC) genes). 632 NLR loci defined by NLR-Annotator correspond to more than one gene in RefSeq v1.0. We looked at these 632 loci in more detail. In 75 cases, we found gaps (stretches of Ns) in the assembly potentially interfering with transcript mapping and thus hampering gene calling and giving rise to two falsely called genes. In 123 cases, we found one of the gene models resembling a complete NLR gene, i.e. the P-loop, at least three consecutive NB-ARC motifs and at least one LRR motif indicating a potential overextension of the NLR locus, usually brought about by a randomly occuring LRR motif shortly downstream of the gene (Table S4). In 28 of 30 random examples (Table S4) of the remaining 434 cases, we observed a stop-codon in the coding sequence interupting the open reading frame in the transcript. In the two remaining cases we were unable to re-construct a consistent gene structure and could not draw conclusions.

We believe that these cases with stop codons represent alleles of NLR genes that have recently been pseudogenized but which are still transcribed. An example supporting this theory is the *Pm2* gene conferring resistance to *Blumeria graminis*, the causal agent of powdery mildew. The *Pm2* gene was cloned from the wheat cultivar Ulka [27] where it encodes a full length NLR. The allele in Chinese Spring has a stretch of 12 nucleotides replaced by five other nucleotides, thus causing a frame shift leading to an early stop codon (Figure S3).

In the subsequent analyses which involve gene models, we proceeded with those genes that resemble a complete NLR gene including the P-loop [59], at least three consecutive motifs associated with the NB-ARC domain (motifs 1, 6, 4, 5, 10, 3, 12, 2) and at least one LRR-associated motif (motifs 9, 11 or 19). Our analyses are thus conducted with a fixed set of 1,540 NLR related genes (Table S5), a number which is most likely an underestimate of the total NLR gene content. Manual curation would likely be required to obtain a more precise estimate of the total true NLR complement.

### Phylogenetic analysis of wheat NLR genes

NLRs constitute a large gene family and only for a very few individual genes has a function been assigned. A phylogenetic analysis provides a means to order this large number of genes with respect to their sequence relationship and arrange them into sub-families. We set out to establish this order and look for common features in various clades of the NLR gene family, which may hint at common functional attributes.

We extracted the NB-ARC domains from NLR protein sequences and calculated their phylogenetic relationships. For this analysis, we used the 1,540 genes rather than loci due to an often-occurring intron within the NB-ARC domain complicating identifiation of the correct, complete open reading frame and subsequent translation without the support of transcript data. Figure 2 shows the NLR phylogeny of Chinese spring with clades highlighted that are discussed in this manuscript. We also marked the closest orthologues of known resistance genes in the tree. Figure 3 shows the same phylogeny but in addition displays the structure of each gene highlighting the conserved amino acid motifs of Coiled coil domain, NB-ARC domain and LRRs with brown, blue and green colors, respectively. Integrated domains are marked with different colors (see respective section).

**Figure 2.**
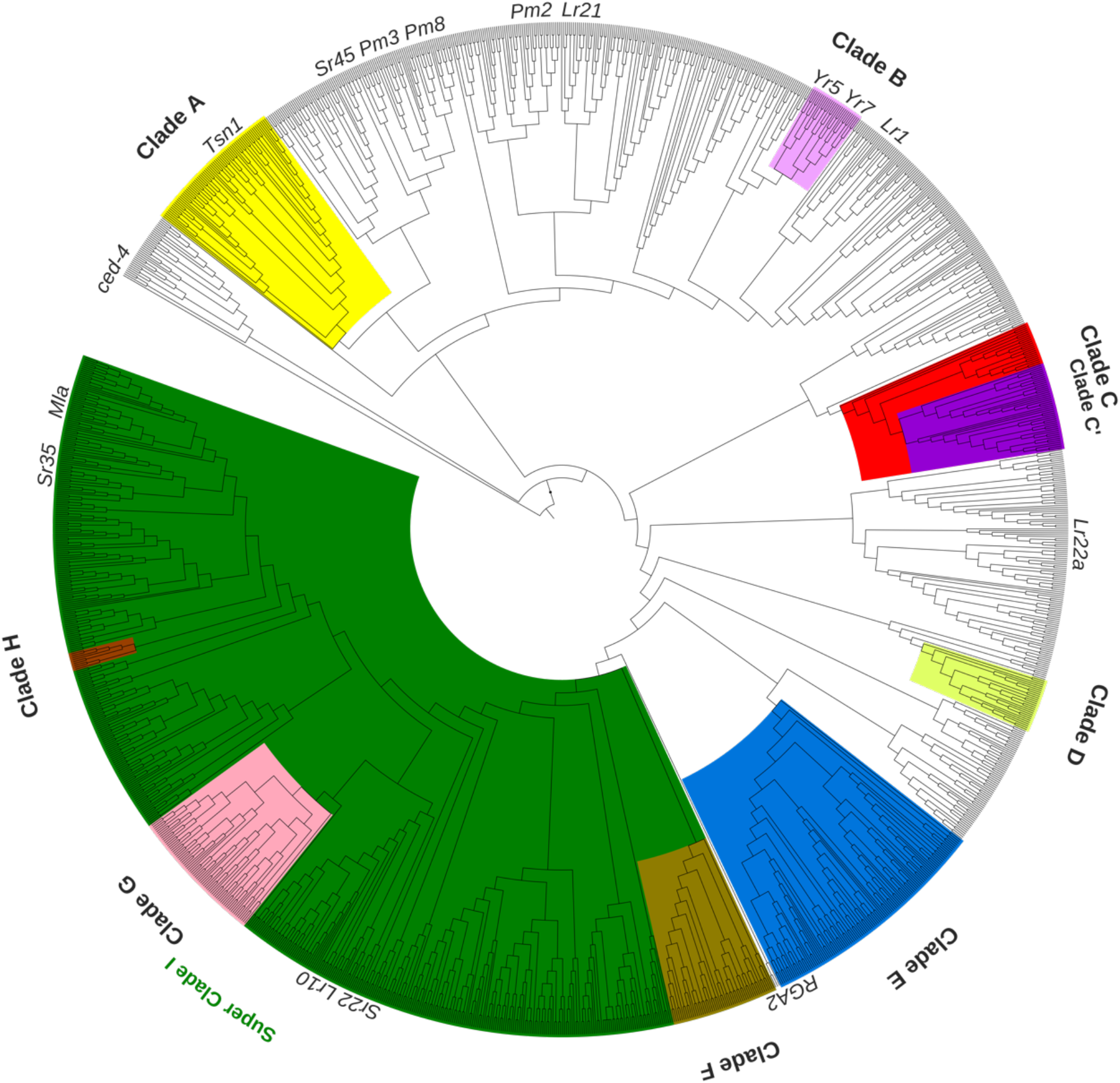
Phylogenetic tree of NLR genes in wheat. Cloned *R* genes are positioned next to their closest homologue in the tree. Specific clades are marked according to certain characteristics: Clades A, C and D as well as Super Clade I contain common intron positions. Clades B (zf-BED), C’ (DDE-TNP), G (various) and H (Jacalin) are enriched for integrated domains. Clade F is enriched for genes hypothesized to be “helper” NLRs. NB-ARC domains of protein sequences from Clade E are preceded by another truncated NB-ARC domain.

**Figure 3.**
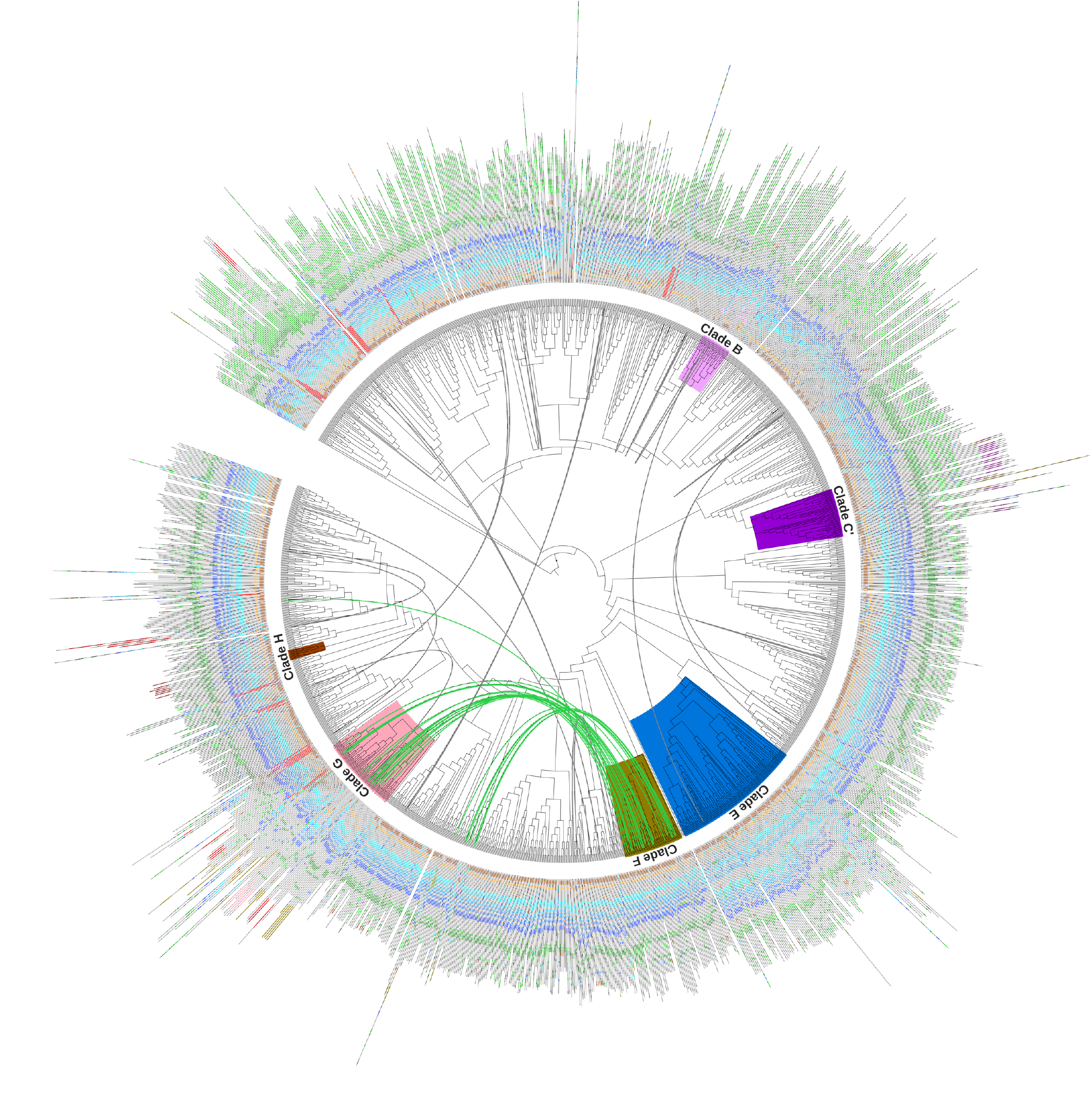
Features of NLR genes in wheat. Connectors (gray/green) between nodes indicates close physical position (<100 kb) of genes and a head-to-head formation. Green connectors mark those where one partner is a member of an orange “helper” clade. The position of motifs within the amino acid sequence are displayed in the outside ring: brown marks CC, blue marks NB-ARC, and green marks LRR motifs. Integrated domains are colour-coded according to: protein kinases (red); DDE (purple); zf-BED (amethyst); kelch (pink); Jacalin (wine).

We also tried to circumvent the problem of not being able to accurately automate the annotation of an NB-ARC domain. To this end we used only the concatenated amino acid motifs within the NB-ARC domain that were identified by NLR-Annotator. The resulting tree (Figure 4) is similar to the tree based on whole NB-ARC protein sequences (Figure 2 and 3). For example, clades with specific features that we identified (see sections below) were mainly preserved in this tree based only on the concatenated NB-ARC motifs. To highlight the similarity between the two trees, we colour-coded NLR loci in the concatenated NB-ARC domain tree corresponding to genes from four distinct clades (denoted B, C’, E and G in Figure 2 for the purpose of this discussion). Apart from eight outliers from Clade G (containing various integrated domains), all loci which clustered in the full-length NB-ARC domain tree (Figure 2), also clustered in the concatenated NB-ARC motif tree (Figure 4). The eight Clade G outliers all contained integrated NB-ARC domains; here, the integrated NB-ARC domain was classified as a separate NLR locus by NLR-Annotator. The other NB-ARC domain of these genes, however, always clustered together with other genes from Clade G as in the full-length NB-ARC domain tree. This shows that if no gene annotation is available, data from NLR-Annotator can directly be used for phylogenetic studies of NLR loci.

**Figure 4.**
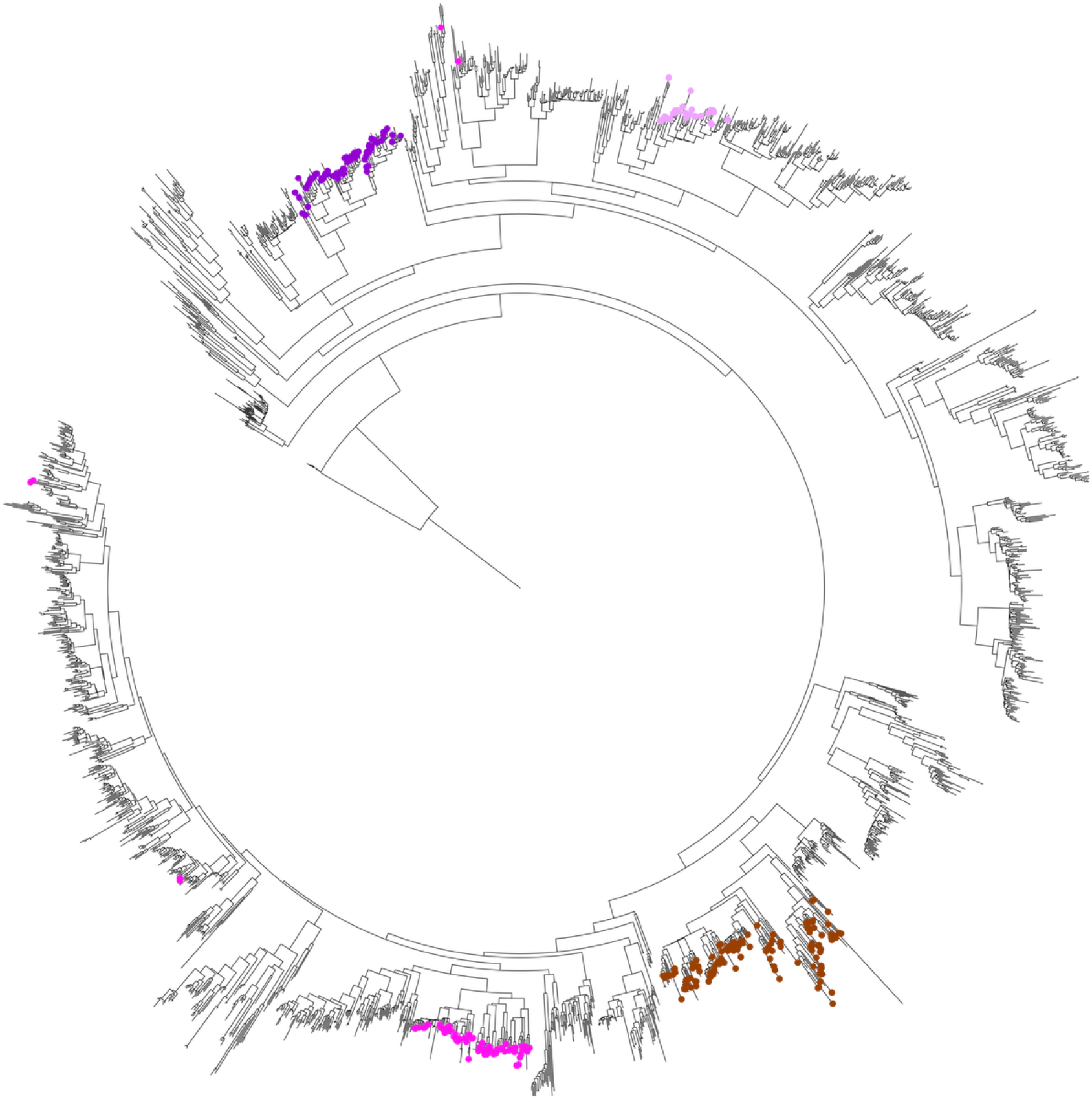
Phylogenetic tree of NLR loci based on concatenated motif sequences of the NB-ARC domain. Colored leaves correspond to loci overlapping with genes in clades (Figure 2) highlighted for integrated DDE domains (purple), zf-BED domains (amethyst), general integrated domains (pink) and elongated NB-ARC domains (brown).

NLRs often appear along the chromosome in physically discrete clusters of sequence-related paralogues. By combining our phylogenetic analysis with the physical location of NLRs we investigated clustering of NLRs in hexaploid wheat both in relation to paralogues on the same chromosome as well as across homeologues on the A, B and D sub-genomes. The most prominent example of an NLR cluster in Triticeae is *Mla*, which has been extensively studied in barley [60–62]. In wheat, the stem rust resistance genes *Sr33*, introgressed from *Aegilops tauschii* and *Sr50*, introgressed from *Secale cereale*, were identified to be homologues of barley *Mla* [31, 33], highlighting the importance of this gene cluster for wheat also. As an example of a complex locus with multiple paralogues, we therefore looked at this region in hexaploid wheat. Phylogenetically, the subclade related to barley *Mla* contains 14 genes on group 1 chromosomes (Figure S4a) or 22 NLR loci when considering the phylogenetic tree based on concatenated motifs (Figure S4b). All wheat *Mla* subclade family members are restricted to small regions (<1.2 Mb) on the short arms of group 1 chromosomes with the exception of the NLR locus chr1A_nlr_75 on chromosome 1A, which is found at a distance of 10 Mb distal of the region (Figure S4b). Pronounced copy number variation of *Mla* homologues, their orientation, and insertion of non-Mla NLRs into the cluster, was observed between the A, B and D subgenomes (Figure S4c). In several cases, complete loci as well as evidence of expression was observed while reading frames were interrupted leading to annotated genes that resemble partial NLRs (e.g. chr1A_nlr16, chr1A_nlr 75 or chr1B_nlr36). In all examples above, the genes had been annotated with high confidence. Locus chr1B_nlr_38 was phylogenetically placed in the clade neighbouring *Mla*. The corresponding annotated gene, TraesCS1B01G035200, however, was placed in the *Mla* clade. This indicates a slight impreciseness in using a mutltiple alignment based on concatenated motifs only in contrast to complete protein sequences for the computation of phylogenetic NLR trees.

### Integrated domains

Some NLR proteins differ from the canonical NLR structure by having additional integrated domains. This structure has been demonstrated in other plant species [17, 18] and has also previously been reported in wheat [63]. It has been proposed that these domains act as a decoy or bait [17, 18], in which the gene from which the integrated sequence originates encodes for a protein that is a target of a pathogen effector(s). A detailed study of additional domains integrated into NLRs can potentially provide information about effector targets in plant pathogen interactions.

We screened the NLR protein sequences for integrated domains. In total, we found 133 of the 1,540 protein sequences carry integrated domains. The most prevalent domains were protein kinases (44 cases), DDE_Tnp_4 (26 cases), and transcription factors (21 cases). Of transcription factors, we found the domains zf-BED, WRKY, AP2, CG-1 and zf-RING. A set of seven genes encode for an NLR with an integrated protein kinase followed by a Motile Sperm domain.

We then associated integrated domains with the phylogeny of the NB-ARC domains of all NLRs. The clades with accumulated integrated domains are shown in Figure 2 and individual domains are displayed within the strucutre of NLRs in Figure 3. In concurrence with previous studies [63, 64], we found that specific clades of the phylogenetic tree are enriched for integrated domains including zf-BED (Clade B, Figure 2), DDE_Tnp_4 (Clade C’, Figure 2) and Jacalin (Clade H, Figure 2). A fourth clade (Clade G, Figure 2) with 69 members is heavily enriched (40 proteins) for various integrated domains: most integrated domains that are not in the abovementioned specific clades (i.e. B, C’ and H) accumulate in this clade (G). Examples include protein kinase-, Kelch-, Grass-, Exo70- and WRKY-domains. The major type of integrated domain that is not exclusively found in a specific clade are the protein kinases.

### Tandem NLRs

NLRs occuring in tandem can function together. They can be arranged in a head-to-head formation in wich the genes are on opposite strands and the distance between gene starts is shorter than the distance between gene ends [65].

We searched for these type of tandem NLR genes in the reference wheat genome. We found 45 NLR pairs with a position as described above and a maximum distance of 50 kb (displayed as internal connectors in Figure 3). Twenty-three of those pairs had one mate located in Clade F. Sixteen of the pairs with one mate located within that clade have the other mate in Clade G, which accumulates various integrated domains. (Figure 2).

### Intron-Exon structure

We investigated whether there was any relationship between the phylogeny and the intron/exon structure of NLR genes. In Super Clade I with 589 members, all but three genes have an intron in the NB-ARC domain after the RNBS-A (motif 6) (Figure 3). Members of this clade include the cloned resistance genes *Lr10, Sr22, Sr33, Sr35* and *Sr50*. The length of the intron varies between 65 and 23,585 bp with an average of 1.7 kb.

Clade E has 65 members and their sequences have a specific variation in the NB-ARC domain consisting of an NB-ARC truncated after the RNBS-B and inserted before a complete NB-ARC domain (Figure S6). This organisation could also be interpreted as a case of an NB-ARC with an integrated domain (see above) consisting of a partial NB-ARC. All genes in this clade also have introns between (i) the attached part and the complete domain, and (ii) within the complete domain between RNBS-C/motif 10 and GLPL/motif 3 (Figure 3). The 69 members of Clade B all have introns before and after the NB-ARC domain. The 64 members of Clade C all have a large intron between RNBS-C/motif 10 and GLPL/motif 3, similar to Clade E but without the characteristic variation in the NB-ARC domain. Finally, the sequences of Clade A (71 members) and Clade D (27 members) have introns between the NB-ARC domain and the LRRs. In summary, we observe specific intron-exon structures within the different clades of the phylogeny computed from NB-ARC domains.

### NLR Expression

In a previous study we observed that in wheat the NLRs encoded by the stem rust resistance genes *Sr22* and *Sr45* were expressed at low levels (∼30/∼60 Gb sequencing resulted in <20x/<15x coverage of *Sr22/Sr45* respectively) in leaves of seedlings [30]. To test whether this observation of low expression could be extended broadly to NLRs in wheat, we sequenced the transcriptomes of young leaves generating more than 300 Gb of data in three samples. We also enriched the same samples for NLR genes (R gene enrichment sequencing; RenSeq [66]) to detect low-abundance NLR transcripts. We considered expression of a transcript detectable if more reads would map to it than necessary to cover the entire length 5-fold. In the combined data sets, we found 1,411 transcripts from 1,124 NLR genes being detectable as transcribed. In the un-enriched transcriptome data, 1104 transcripts (78.2%) from 921 genes could be detected. Down-calculating those numbers (Figure 5), we estimate that generating 50 Gb of wheat transcriptome data would allow for 50% of NLRs expressed in young leaves to be detected.

**Figure 5.**
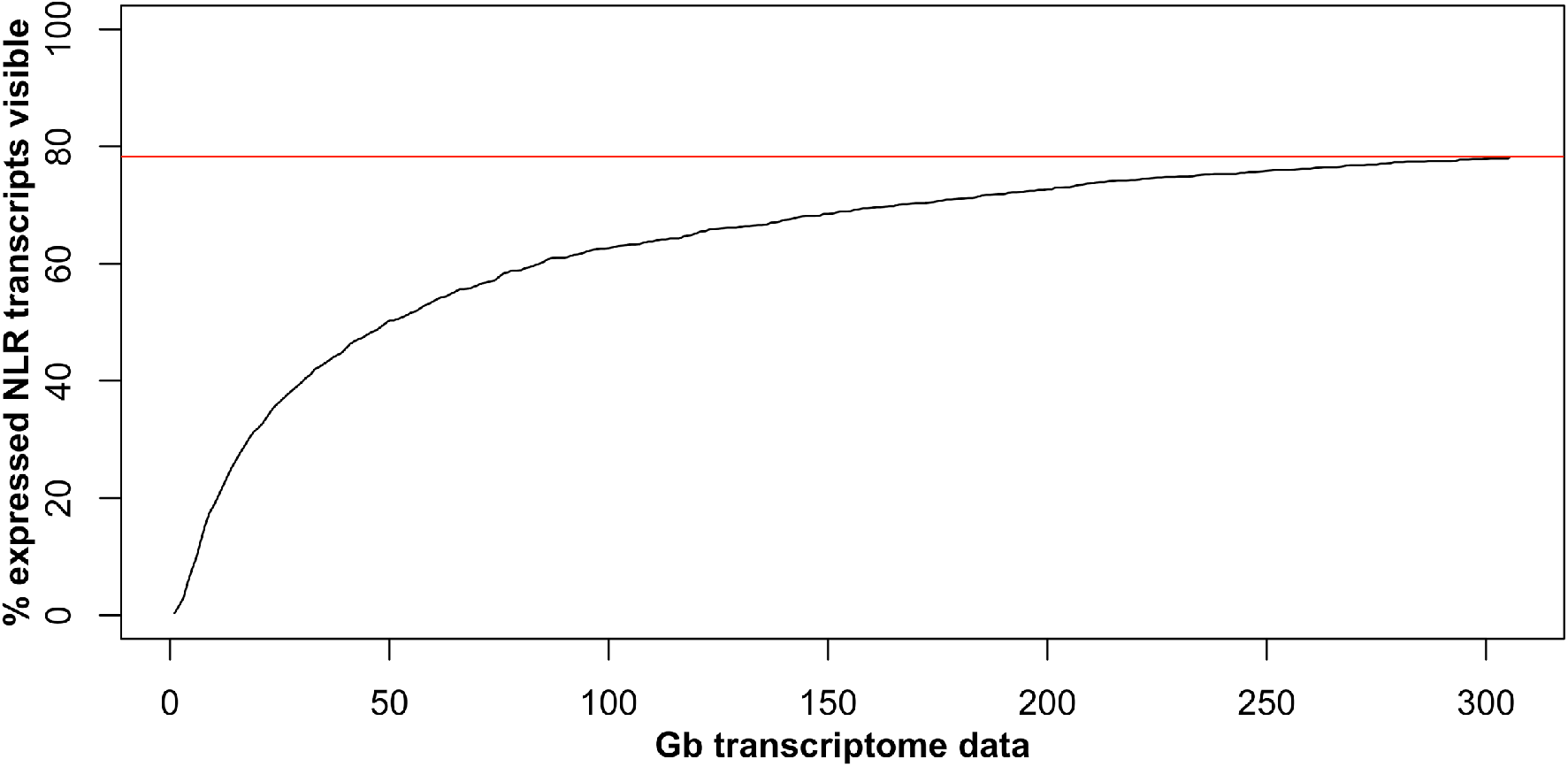
Estimated detectability of NLR transcripts in transcriptome studies. The percentage of expressed NLR transcripts (Y-axis) in leaves from 3-leaf stage are shown in relation to the amount of sequencing data (X-axis) used as an input. A transcript is considered detectable if the combined length of mapped reads exceeds 5 times the length of the transcript. The red line indicates the maximum percentage of NLR transcripts detected with 305 Gb of RNA-Seq data.

### Developmental and stress-induced expression of NLRs

There are more than a thousand NLRs in a wheat genome, representing 1-2% of the total gene coding capacity [35]. However, the immunity conferred by NLRs comes at a metabolic cost and over-activation of immune responses may give rise to stunted growth [67, 68]. Therefore, we assume that NLR expression would be tightly regulated during development or under stress.

We investigated how NLRs are expressed in different tissues, development stages and in response to biotic stress. We used two publicly available data sets containing transcriptomes for multiple tissues and development stages [40, 68] and also generated our own data set using RenSeq on cDNA. The latter included cDNA isolated from roots, seedling leaves, mature leaves, flag leaves, stems, heads and seed. The enrichment procedure may introduce a bias depending on enrichment efficiency. Comparison of data from total RNAseq and RenSeq cDNA of young leaves revealed a (median) 506-fold increase in NLR-associated transcripts in the RenSeq data (Figure S6). Although there was a good general correlation between the two datasets (Pearson r-value 0.7, p-value < 1e-16), the bias introduced by enrichment was significant, indicative of a certain limitation when using the cDNA RenSeq data to study gene expression. Notwithstanding, hierarchical clustering of the different cDNA RenSeq tissue samples showed that the biological replicates clustered together, while the different tissue samples and developmental stages were clearly separated (Fig. S7 a). Using the two public RNAseq datasets, which we mined for NLRs, we observed a similar trend in the separation of different tisssue samples (Figure S7 b and c). This indicates that the bias introduced by cDNA RenSeq is sufficiently consistent in all experiments to allow us to proceed to compare NLR expression across tissues and developmental stages. The clearest distinction was observed between the root and other samples, while the samples from the 3^rd^ and 7^th^ leaf stage clustered close together or overlapped. On the other hand, the flag leaf samples were clearly distinct from the 3^rd^ and 7^th^ leaf stage samples, which, surprisingly clustered more closely with the stem and head samples.

We also checked whether NLR expression changes after biotic stress. Plants also respond to conserved pathogen or microbe associated patterns (PAMPs or MAMPs) such as chitin, a constituent of fungal cell walls, or flagellin from the bacterial flagellum. Molecular events that occur during PAMP-triggered immunity (PTI) are highly conserved between dicots and monocots and include rapid production of reactive oxygen species (ROS), MAP kinase induction and defence gene induction [69]. We hypothesised, that as a first layer of defence, the PAMP triggered immune response might also include increased expression of NLRs to help prime the plant defence system for effector triggered immunity. To test this hypothesis, we infiltrated young wheat plants (at the three leaf stage) with two PAMPs, namely chitin [70] and flg22, a 22 amino acid peptide from flagellin [71], and then sequenced the transcriptomes of water treated and PAMP-challenged plants at 30 minutes and 3 hours after challenge. Three (non-NLR) genes that had previously been reported to be PAMP responsive showed differential expression [72] indicating a successful PAMP challenge (Fig. S8).

We found that 54 NLR genes for flg22 and 79 NLR genes for chitin were strongly up-regulated 30 minutes after challenge whereas expression levels at 3 hours post challenge were not different from the water treated samples (Figure S9). The upregulated NLR genes were not associated with specific phylogenetic clades.

## Discussion

In this study we present the tool NLR-Annotator for *de novo* annotation of NLR loci. Our tool distinguishes itself from standard gene annotation in that it does not require transcript support to identify NLR-associated loci. NLR-Annotator is therefore a powerful tool for the identification of potential *R* genes that could be used in breeding for resistance to wheat pathogenes. Very often, however, the funcional copy of an *R* gene is not present in the accession that was chosen for a reference genome. The positive selection imposed on NLRs often results in extensive accessional sequence and copy number variation at NLR loci [19–22]. Thus, the corresponding NLR homologue in the wheat reference accession, Chinese Spring, would not be considered by a standard gene annotation. NLR-Annotator, on the other hand, is not limited to functional genes. In our analysis of Chinese Spring, we found 3,400 loci while only 1,540 complete NLRs could be confirmed through gene annotation in IWGSC RefSeq v1.0. Within the additional 2,360 loci identified by NLR-Annotator there might be some NLRs with potential for function. However, we speculate that most of these are non-functional pseudogenes. Nevertheless, it is important to define these loci in the reference accession Chinese Spring, because these genes may have functional orthologues in other accessions. A case in hand concerns *Pm2* (from the cultivar Ulka), which in Chinese Spring has an out-of-frame indel leading to an early stop codon. The corresponding locus in Chinese Spring is still transcribed at a detectable level. The frameshift interrupting the open reading frame led to two remaining still large enough open reading frames, which then have been annotated as separate genes.

In some cases (e.g. transcript studies or inspection of integrated domains), an analysis of NLRs requires functional genes. Previous work on describing NLRs in earlier versions of the Chinese Spring genome, have searched within the genome-wide gene annotation for NLRs. This led to fewer NLRs being identified, for example 16 full-length NLRs in the 2014 Chinese Spring Survey sequence [38, 40] and 1,174 full-length NLRs in the 2017 Chinese Spring W2RAP assembly [40]. For our present study, we also improved the general annotation by supplying it with cDNA data enriched for NLR transcripts through NLR enrichment sequencing, and secondly, we developed NLR annotator to identify NLRs independently of gene annotation.

To improve the annotation further would require manual annotation locus-by-locus, in each case, taking into account existing gene annotations, mapped transcript data from different sources, and our NLR-Annotator loci. All these data would have to be integrated and inspected to provide *bona fide* gene models. Given the diversity of NLR complements in different wheat accessions [26] the amount of work required for this effort is considerable. For practical applications such as *R* gene cloning, our automated NLR locus discovery will in most cases likely be sufficient to identify a candidate gene, which can be annotated *ad hoc* in the relevant accessions. For studies requiring annotated genes we have to settle for a subset of the actual NLR complement.

We set out to investigate the sequence diversity relationships within the NLR gene family in Chinese Spring. It has been well documented that in NLRs the NB-ARC domain is far more conserved than the LRRs. Therefore, phylogenetic studies compare the NB-ARC domains to explore and depict the sequence relationship between NLRs [46, 48, 63]. Extracting NLR domains from NLR loci defined by NLR-Annotator is complicated by the presence of introns within domains the boundaries of which can only be defined properly using transcript data. In addition, we were interested in cataloging the occurance and distribution of integrated domains in the NLRs. This can only done on annotated genes. Therefore, in our subsequent analysis, we used the 1,540 annotated NLRs to compute a phylogenetic tree.

With the reference sequence, we studied the phylogeny, integrated domains, and helper NLRs. This has been recently described in detail by Bailey and colleagues [63] based on analysis of the W2RAP assembly of wheat. In brief, we found that the new assembly supports the observations made in the study by Bailey and colleagues. One additional feature that we discovered is a clade of NLRs which contain elongated NB-ARC domains (Clade E, Figure 2). The elongated part of the NB-ARC domain precedes the canonical NB-ARC domain and consists predominantly of motifs 6, 4 and 5, often preceded by motif 1 (See Figures S1 and S5). We do not know the function of this elongated NB-ARC domain. However, a member of this clade is RGA2 which has previously been described to be essencial for Lr10 function [56].

To expand the previous phylogenetic analysis to include the entire set of all 3,400 NLR loci, we developed a new feature in NLR-Annotator. By reducing the NB-ARC domains to only be represented by the even more conserved motifs (Supplementary Figure 1), we cannot only ensure to exclude introns from the phylogenetic analyses, but we can also exploit the *a priori* knowledge of the position of the motifs within the NB-ARC domain to avoid an external multiple alignment. This then permits printing out the alignment from NLR-Annotator to be used directly as an input to compute a phylogenetic tree. The resulting phylogeny (Figure 4) was validated by visualising leaves that correspond to genes from specific clades in the phylogeny derived from gene models (Figure 2). The phylogenetic relationship of genes was maintained for corresponding loci in the tree generated from NLR-Annotator. One example, however was found in the *Mla* region, where a locus was associated with a neighboring sub-clade rather than the other loci of the *Mla* region.

Finally, with the IWGSC RefSeq v1.0 gene annotation v1.0, we performed NLR expression profiling. We found that many NLRs are expressed at low levels, requiring either extremely deep sequencing or target enrichment to be detected. In addition, the expression of many NLRs was tissue specific. In particular, roots had a very different profile from the aerial tissues (e.g. leaves, stem and head). We speculate this difference may reflect the specifity of certain NLRs towards tissue specific diseases. For example, an NLR guarding a celluar hub specifically expressed in roots, does not need to be expressed in leaves. We also looked at NLR induction in response to PAMPs and found that a subset of NLRs are triggered following PAMP exposure. The transcriptional response was measurable within 30 minutes and had returned to base level after 3 hours. This response shows the tight regulation of the NLR immune system and may reflect the fine tuning between the cost of defence and resource allocation for growth and reproduction.

## Conclusions

In this study, we demonstrate the power of the new reference sequence of wheat cultivar Chinese Spring in allowing a comprehensive cataloging and positioning of a complex multigene family – the NLRs. The analysis of NLRs was further faciliated by the development of NLR-Annotator, a tool which allows the automated annotation of NLRs across a whole genome. We predict this will facilitate the cloning of functional *R* genes from non-reference accessions of wheat, by positioning NLR loci within mapped intervals, as demonstrated for leaf and stem rust, two major diseases of wheat.

## Methods

### NLR Annotator

The NLR Annotator pipeline is divided into three steps: (1) dissection of genomic input sequence into overlapping fragments; (2) NLR-Parser, which creates an xml-based interface file; (3) NLR-Annotator, which uses the xml file as input, annotates NLR loci and generates output files based on coordinates and orientation of the initial input genomic sequence. All three programs are implemented in Java 1.5. Source code, executable jar files and further documentation has been published on GitHub (https://github.com/steuernb/NLR-Annotator). The version of NLR-Parser used here has been published previously along with the MutRenSeq [30] pipeline and uses the MEME suite [73]. All genomes annotated in this publication were dissected into fragments of 20 kb length overlapping by 5 kb. Manual investigation of questionable *A. thaliana* protein sequences was performed using SMART [74].

### *R* gene intervals

Coding sequences of cloned *R* genes were aligned to pseudo chromosomes of Chinese Spring using BLASTn [75] and default parameters. To align marker sequences extracted from the literature to the pseudo chromosomes we used BLASTn and the parameter –task blastn to amend for short input sequences. In uncertain cases, matching positions were validated using DOTTER [76].

### Selection of NLR genes from gene annotation

To identify NLRs in the RefSeq v1.0 gene annotation, NLR-Parser was run on CDS of gene models. Tab-separated output from NLR-Parser was generated using output option –o. The last column in that table is a list of identified motifs [48]. Genes were selected by presence of motif 1 (p-loop), presence of either motif 9, 11 or 19 (LRR motifs) and at least one of the following consecutive motif combinations: 1, 6, 4; 6, 4, 5; 4, 5, 10; 5, 10, 3; 10, 3, 12; or 3, 12, 2. Motifs detected in reverse frames of sequences were discarded (One case; TraesCS3D01G521000LC). Examples from cases with discrepancies between NLR-Annotator and gene annotation were manually inspected using Web Apollo [77].

### Phylogenetic analysis

Protein sequences from NLR genes were screened for motifs associated with NB-ARC domains using MAST [73]. Intervals within each sequence were defined based on presence of motifs 1, 6, 4, 5, 10, 3, 12, 2. An interval has to start with motif 1, other motifs may be absent but if present, the order is not allowed to be changed. For each protein sequence the largest interval including 20 flanking amino acids was used as NB-ARC domain. A multiple alignment of NB-ARC domain sequences was generated using clustalw2 [78]. A phylogenetic tree was generated using FastTree [79]. The tree was visualized using iTOL [80].

### Integrated domains analysis

Protein sequences of NLRs were screened for domains using HMMER v. 3.1b1 [81] and PFAM-A v. 27.0 (http://pfam.xfam.org/). Domains with an e-value less than 1E-5 were regarded and filtered for domains usually overlapping with the NB-ARC domain (NACHT, AAA) as well as the LRR domain. Java source code was deposited on github (www.github.com/steuernb/wheat_nlrs)

### Tandem NLR analysis

General feature format (GFF) files of both high- and low-confidence genes from the IWGSC RefSeq annotation v1.0 were screened for pairs of NLR genes that were on reverse complementary strands, in a distance of less than 50 Kb and in a head-to-head relation, i.e. the distance between gene starts is shorter than the distance between gene end. Java source code was deposited on github (www.github.com/steuernb/NLR_manuscript)

### Growing Chinese Spring and extracting RNA

Plants of cultivar Chinese Spring (CS42, accession Dv418) were grown in the green house. The tissues were taken from different parts of the plants at various growth stages. The total RNA was extracted with RNeasy Plant Mini Kit (QIAGEN Cat. No 74904) following the company’s protocol. The total RNA samples were then digested with DNAse I (11284932001 Roche) using the company’s protocol. After the DNA digestion, the total RNA samples were purified with Agencourt AMPure XP beads (Agencourt part number: A63881) at 0.5x rate.

### Elicitation with PAMPs

The protocol was modified from Schoonbeek et al. [72]. Chinese Spring wheat plants were grown for 3 weeks in a growth cabinet under a 16:8 hours day:night regime at 23:18 °C. For each biological repetition three strips (2 cm) where cut from leaf 2 and 3, placed in a 2 ml tube with sterile water and vacuum-infiltrated for 3 times for 1 minute. The following day water was removed and replaced by fresh water or PAMPs dissolved in water at 1 g/l for chitin (Nacosy, YSK, Japan) or 500 nM flg22 (www.peptron.com). Samples were drained and flash frozen in liquid Nitrogen after 30 or 180 min prior to pulverisation with 2 stainless steel balls in a Geno/Grinder (SPEX). RNA was extracted using the RNAeasy plant kit (www.Qiagen.com), the concentration determined on a NanoDrop 8000 Spectrophotometer (Thermo Fisher Scientific) and quality assessed with a RNA 6000 Nano chip on a Bioanalyzer 2100 (Agilent Technologies). After removal of genomic DNA with TURBO DNA-free™ Kit (Thermo Fisher Scientific), 1 μg of RNA was converted to cDNA with SuperScript IV (Thermo Fisher Scientific) and expression of PAMP-inducible genes [72] verified by quantitative RT-PCR (Figure S8) using 0.4 μl of cDNA per 16 μl reaction with PCR SYBR Green JumpStart Taq ReadyMix (Sigma) on a LightCycler 480 (Roche Life Science). Gene-expression of 3 biological replicates is expressed as log2 relative to the expression level at t=0, after infiltration and overnight incubation but before addition of fresh water or PAMP-solutions. Expression was normalised to EF-1α (Elongation factor 1-alpha, M90077; with primers ATGATTCCCACCAAGCCCAT and ACACCAACAGCCACAGTTTGC). Genes tested were syntaxin (homologous to barley ROR2, tplb0005g03; TCGTGCTCAAGAACACCAAC and AATCGAGTGGCTCAACGAAC), EFE (Homologous to parsley EFE (Immediate-early fungal elicitor protein CMPG1 and pub23 (Homologous to barley and arabidopsis E3 ubiquitin-protein ligase PUB23, Traes_3B_0B86BCF93; CGTTCATCAGAATGCTCAGCTG and TTCTCTTTTGTAGGCACGAACCA).

### Enrichment of cDNA

cDNA for targeted enrichment was prepared with KAPA Stranded mRNA-Seq Kit (F. Hoffmann-La Roche AG, Basel, Switzerland) following the manufacturer’s protocol with minor modifications. Briefly, 5 μg of total RNA was used to capture mRNA with magnetic oligo-dT beads, followed by fragmentation for 6 min at 85 °C to generate 300-400 nt fragment sizes. First and second strands were synthesized according to the manufacturer’s protocol. No A-tailing step was performed and adapter (100 nM final concentration) from NEBNext Ultra DNA Library Prep Kit for Illumina (New England Biolabs, Inc., Ipswich, MA, USA) was used for ligation. After ligation, 3 μl of USER enzyme (New England Biolabs) were added to the reaction and incubated at 37 °C for 30 minutes. Post ligation purifications and amplification (8 cycles) were performed according to the manufacturer’s protocol only substituting KAPA Library Amplification Primer Mix (10X) with NEBNext Universal PCR Primer for Illumina and NEBNext Index primers for Illumina (2 μM final concentrations each). Amplified libraries were purified using Agencourt AMPure XP beads (Beckman Coulter, CA, USA) with a 1:0.65 ratio of amplified cDNA to beads. Individual libraries were quantified with Quant-iT PicoGreen dsDNA Assay Kit (Thermo Fisher Scientific, Waltham, MA, USA) and equimolar amount used for targeted enrichment with custom MYcroarray MYbaits kit (Arbor Biosciences, MI, USA) and the corresponding protocol. Enriched libraries were sequenced on an Illumina HiSeq 2500. Raw data was deposited at SRA under study ID PRJEB23081.

### Expression data analysis

The transcript analysis pipeline (calculation of read counts and TPM (transcripts per million)), is based on Kallisto [82], and is common with the global wheat study by Ramírez-González *et al*. [42]. Complete transcript lists were filtered for NLR genes as defined above. To estimate the detectability of NLR transcripts, we combined Kallisto read counts from 3-leaf stage replicates. We considered a transcript to be expressed if either in total RNA or in RenSeq cDNA the combined length of mapped reads exceeded 5 times the length of the transcript itself. Testing the same criterion using only read counts based on 305 Gb of total RNA input data, we found 78% of NLRs. We then gradually reduced Kallisto read counts proportionally to a reduced input data to estimate the percentage of NLRs detected with less input data.

Hierarchical clustering of samples was performed using R. A Pearson correlation was used as distance function. R (https://www.r-project.org/) was used for calculation and plotting.

## Declarations

### Availability of data and material

The datasets generated and/or analysed during the current study are available in the EBI short read archive (SRA) under study numbers PRJEB23081 and PRJEB23056.

### Author contributions

General concept and design of studies: BS, KW, JJ, BK, SK, BW. Design and testing of NLR-Annotator: BS, KW. Resistance gene mapping to RefSeq: BS, SK. Analysis of integrated domains and NLR pairs: BS, EB, KK. Experimental concept and design of tissue-specific RNA-Seq: BS, KW, GY, CU, BW. Experimental concept and design of PAMP-triggered NLR-Expression: BS, HS, CR, BW. Tissue collection and RNA extraction: GY. cDNA RenSeq: KW, AW. Bioinformatics analysis and figures: BS, RR, IY. Manuscript draft: BS, BW. Manuscript review: All authors read and approved the final manuscript.

## Acknowledgements

We thank the IWGSC for early access to the RefSeq v1.0 of Chinese Spring, our colleagues Yajuan Yue and JIC Horticultural Services for plant husbandry, and the NBI Computing Infrastructure for Science (CiS) group for HPC maintenance. We thank David Swarbreck and Gemy Kaithakottil for technical support with Web Apollo.

## Supplementary Figure Legends

**Figure S1.**
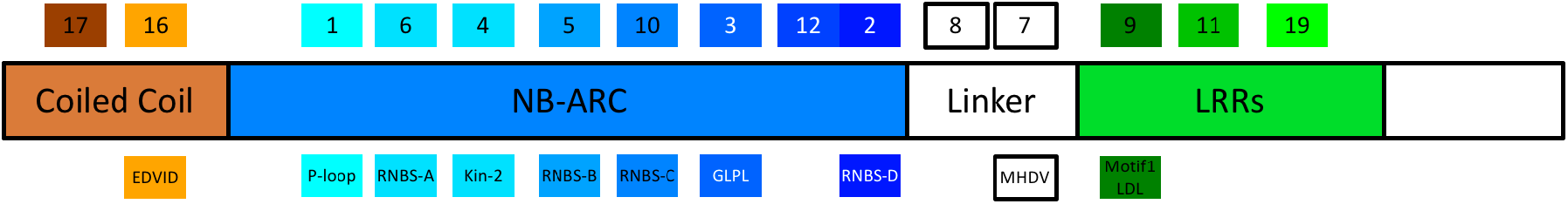
Consensus Structure of a CC-NLR. Numbered rectangles on top depict conserved amino acid motifs within NLRs published by Jupe and colleagues [48]. Rectangles below are conserved motifs identified by Meyers and colleagues [59]. LRR-associated motifs usually occur in varying order and copy number.

**Figure S2.**
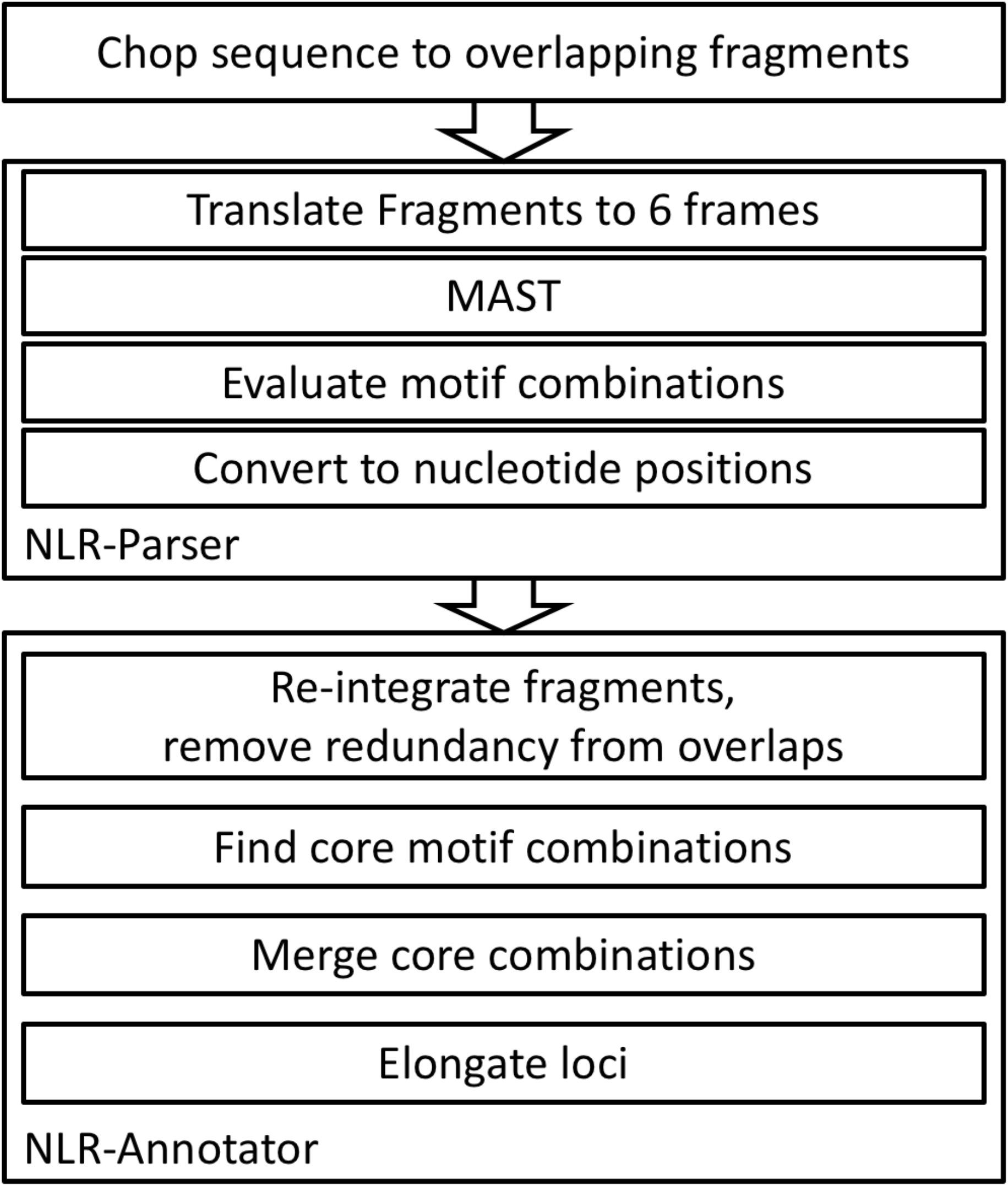
Workflow of the NLR-Annotator pipeline.

**Figure S3.**
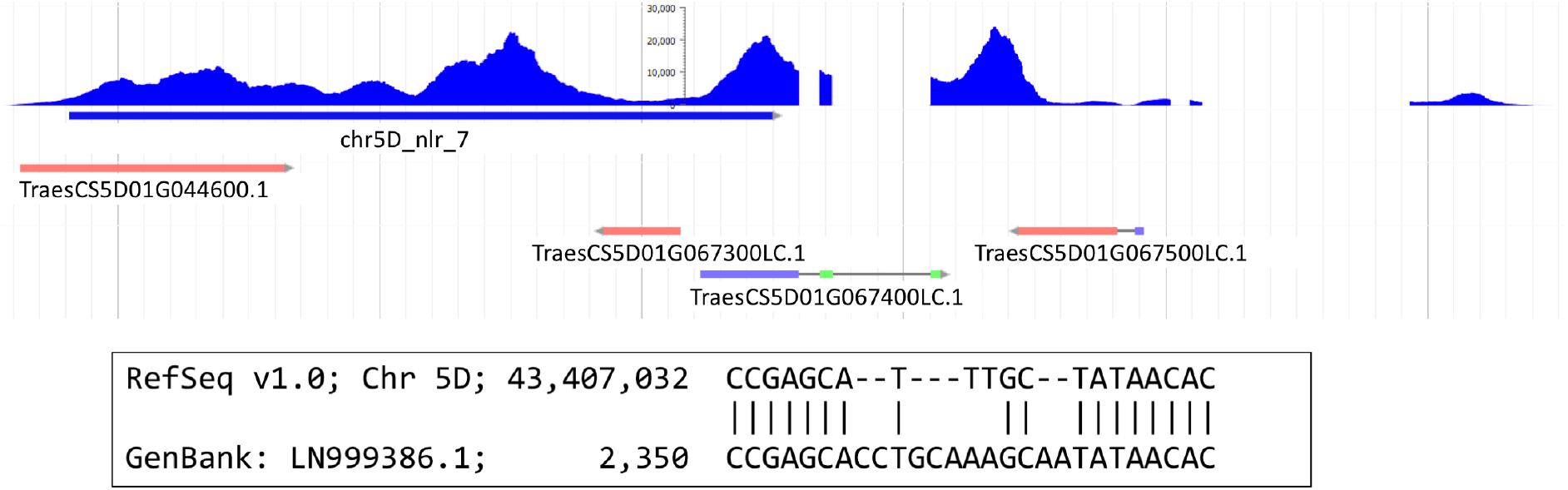
Example of problems with NLR gene annotation in RefSeq v1.0. The blue histogram on top visualizes transcript support from NLR-enriched cDNA. The blue bar below is an NLR locus called with NLR-Annotator. Below are gene models from IWGSC RefSeq v1.0 annotation that overlap with the NLR locus. This region is the best hit for the *Pm2* resistance gene [27]. Twelve bases in the original coding sequence of *Pm2* have been replaced by five bases in Chinese Spring causing a frame shift and an early stop codon (boxed insert).

**Figure S4.**
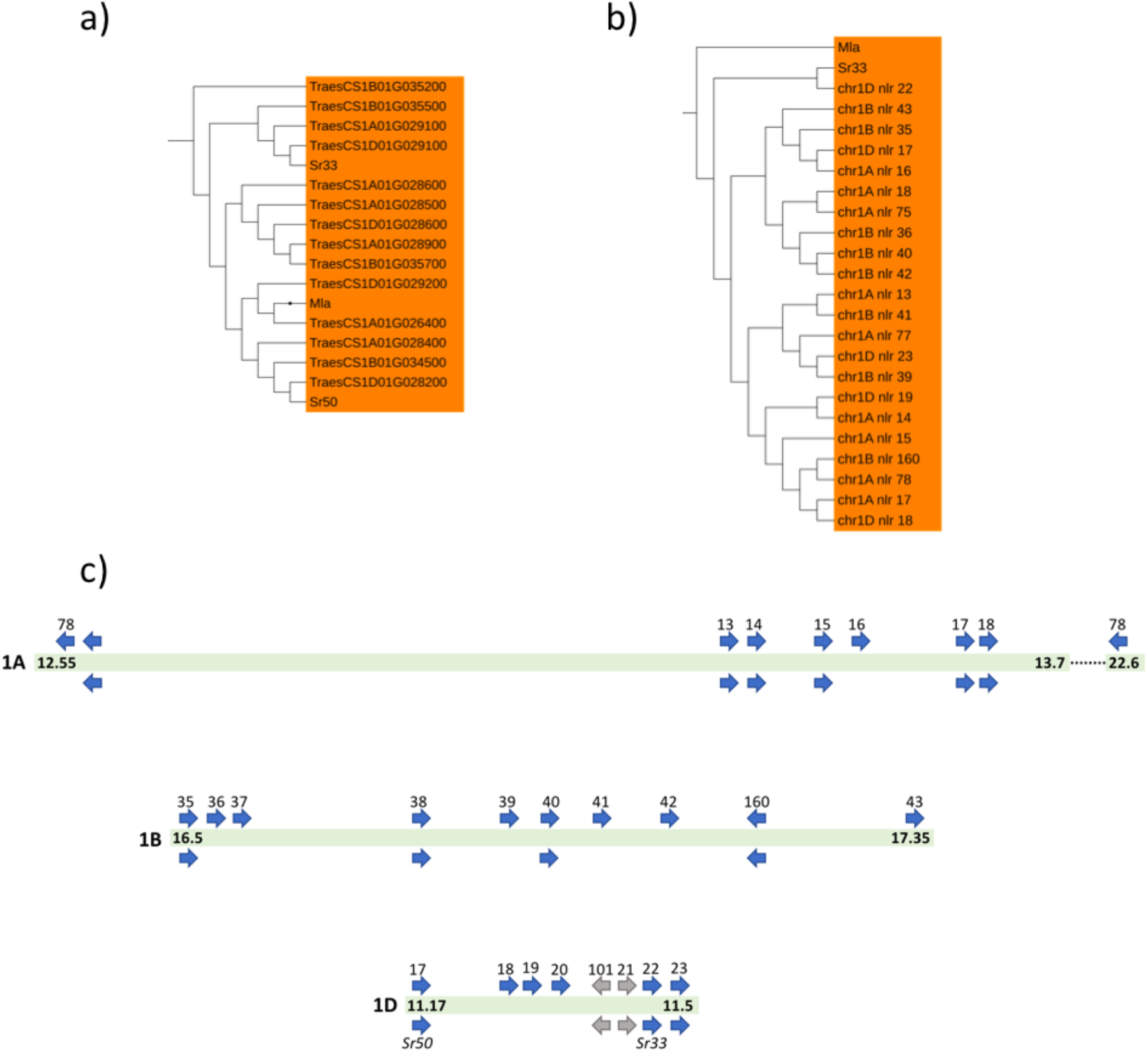
The *Mla* region in Chinese Spring. a) shows the IWGSC RefSeq v1.0 genes that are members of the Mla containing sub-clade in the NLR phylogeny. *Sr33, Sr50* and *Mla* have been added as well. b) shows the corresponding sub-clade of loci predicted with NLR-Annotator. c) shows the Wheat group 1 chromosome regions orthogonal to the barley *Mla* region. Numbers on green bars indicate start and end (in Mb) on group 1 chromosomes. Arrows above bars indicate loci as found by NLR-Annotator. Arrows below indicate annotated gene models in RefSeq v1.0. Arrows in blue indicate close homology to *Mla*.

**Figure S5:**
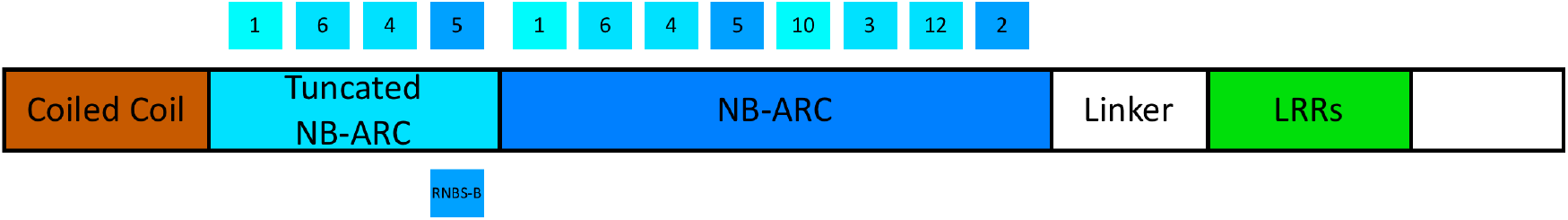
Consensus structure of NLRs from Clade E (see Fig. 2).

**Figure S6.**
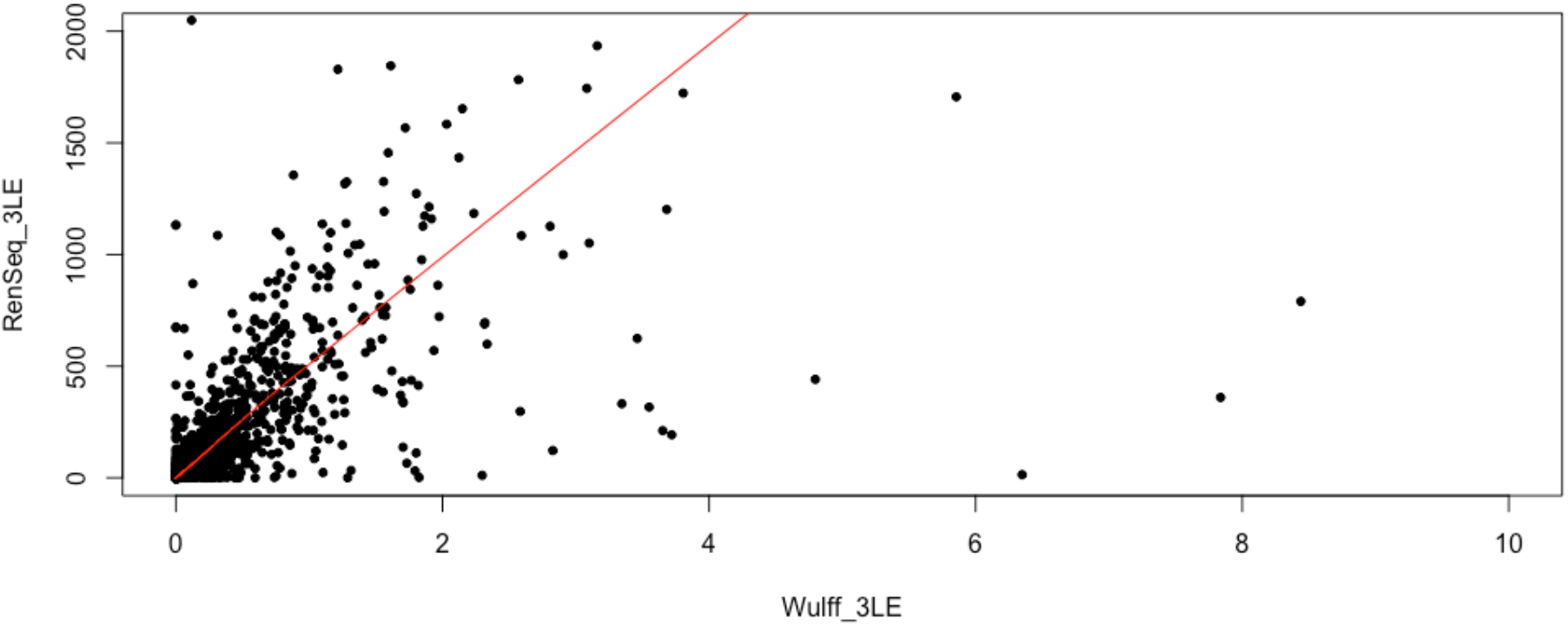
Scatter plot of Transcript Per Million (TPM) values of RenSeq cDNA and total RNA-Seq. Red line indicates lowess-fitted regression.

**Figure S7.**
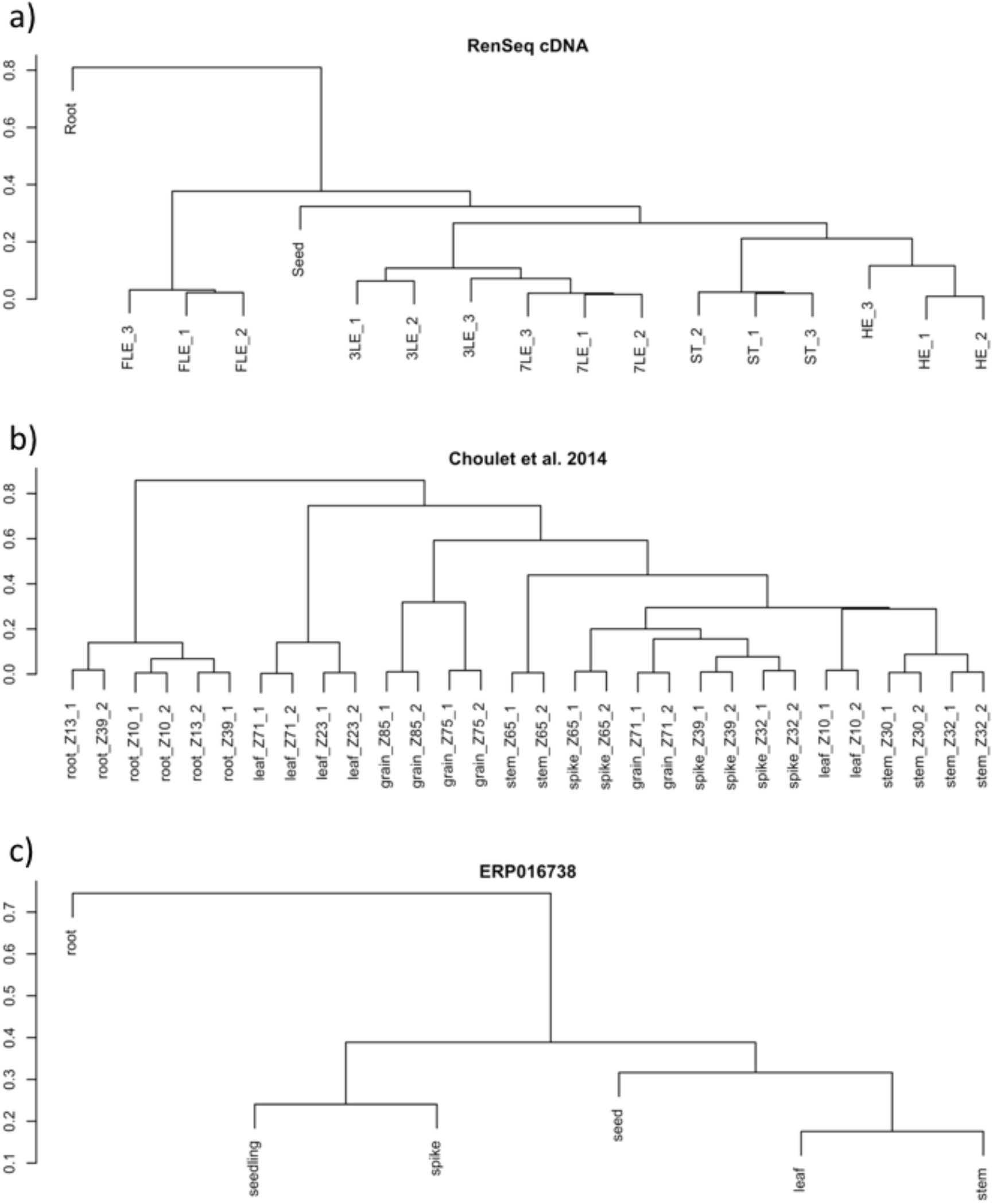
Hierarchical clustering of gene expression in different tissues. Clustering is based on Pearson correllation of TPM values of NLR genes.

**Figure S8.**
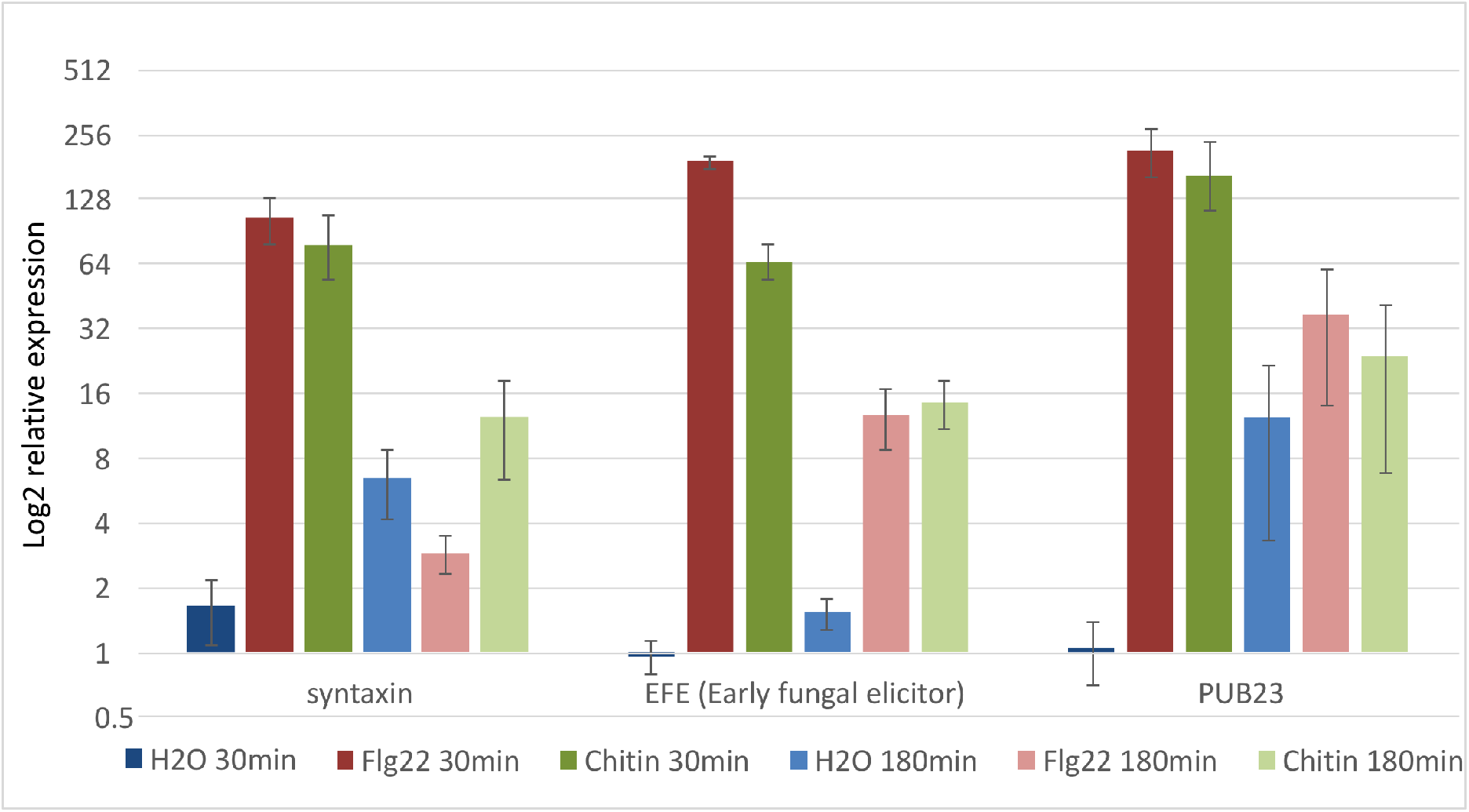
Expression of known PAMP inducible Genes in RNA samples. Expression was measured using quantitative RT-PCR

**Figure S9.**
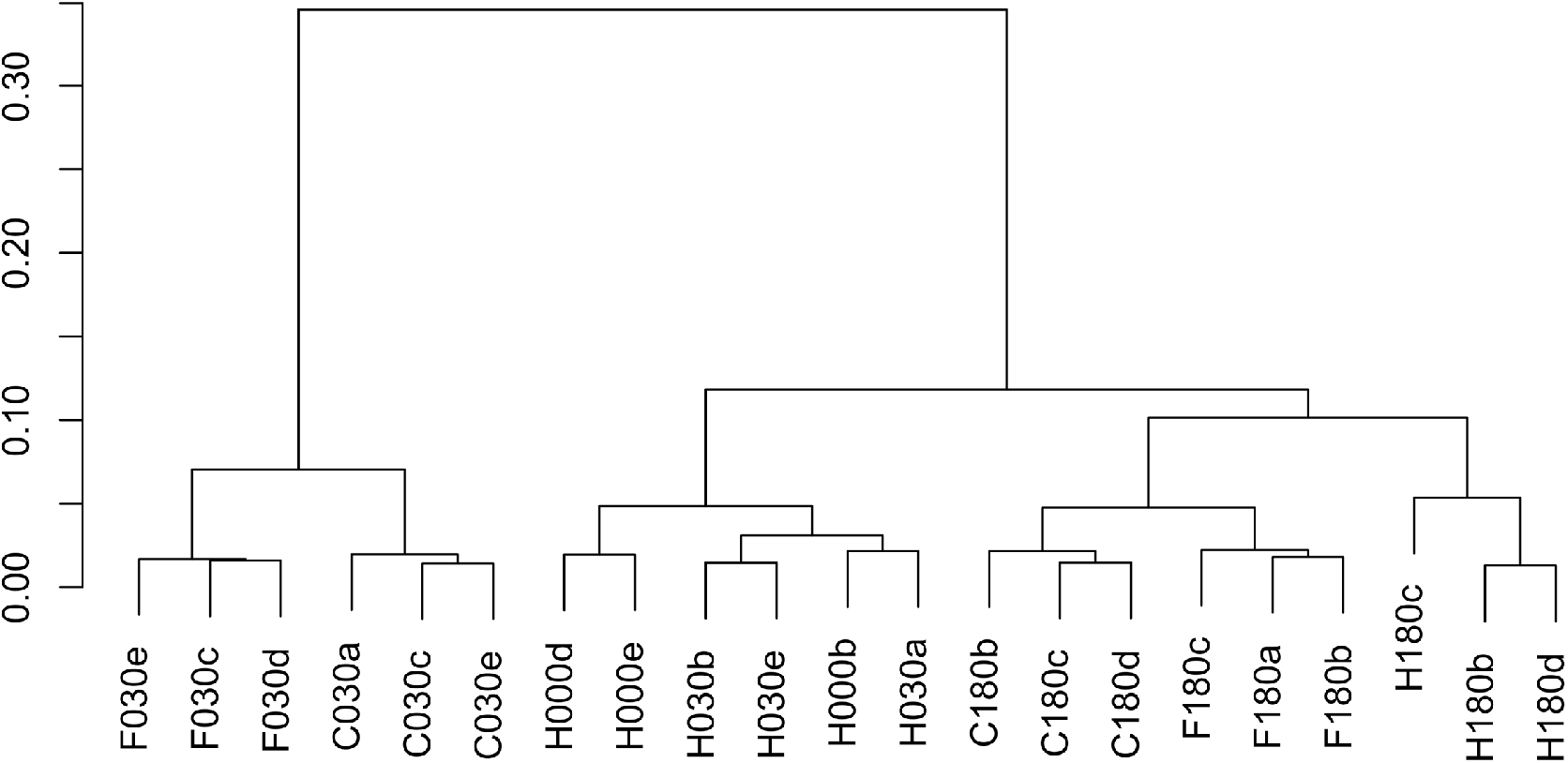
Hierarchical clustering of gene expression in leaves from wheat plants challenged with flagellin and chitin.

### Supplementary Tables

Table S1. Number of NLR loci found in exemplary plant genomes.

Table S2. Number of NLR loci per wheat chromosome.

Table S3. Candidate NLR loci in *Sr* and *Lr* disease resistance gene map intervals projected onto Chinese Spring.

Table S4. NLR loci and overlapping genes from RefSeq annotation v1.0.

Table S5. List of genes that were confirmed to be clomplete NLRs.

Table S6. Domains found in NLR genes.

Table S7. TPM values for NLR genes under biotic stress condition.

## References

1. FAOSTAT [http://www.fao.org/faostat/]

2. Pretorius ZA, Singh RP, Wagoire WW, Payne TS: Detection of Virulence to Wheat Stem Rust Resistance Gene *Sr31* in *Puccinia graminis*. f. sp. *tritici* in Uganda. Plant Dis 2000, 84(2):203–203.

3. Olivera P, Newcomb M, Szabo LJ, Rouse M, Johnson J, Gale S, Luster DG, Hodson D, Cox JA, Burgin L et al: Phenotypic and Genotypic Characterization of Race TKTTF of *Puccinia graminis* f. sp. *tritici* that caused a Wheat Stem Rust Epidemic in Southern Ethiopia in 2013-14. Phytopathology 2015, 105(7):917–928.

4. Shamanin V, Salina E, Wanyera R, Zelenskiy Y, Olivera P, Morgounov A: Genetic diversity of spring wheat from Kazakhstan and Russia for resistance to stem rust Ug99. Euphytica 2016, 212(2):287–296.

5. Firpo PDO, Newcomb M, Flath K, Sommerfeldt-Impe N, Szabo LJ, Carter M, Luster DG, Jin Y: Characterization of *Puccinia graminis* f. sp *tritici* isolates derived from an unusual wheat stem rust outbreak in Germany in 2013. Plant Pathol 2017, 66(8):1258–1266.

6. Bhattacharya S: Deadly new wheat disease threatens Europe’s crops. Nature 2017, 542(7640):145–146.

7. Islam MT, Croll D, Gladieux P, Soanes DM, Persoons A, Bhattacharjee P, Hossain MS, Gupta DR, Rahman MM, Mahboob MG et al: Emergence of wheat blast in Bangladesh was caused by a South American lineage of *Magnaporthe oryzae*. BMC Biol 2016, 14.

8. Abu Sadat M, Choi J: Wheat Blast: A New Fungal Inhabitant to Bangladesh Threatening World Wheat Production. Plant Pathology J 2017, 33(2):103–108.

9. Fraaije BA, Bayon C, Atkins S, Cools HJ, Lucas JA, Fraaije MW: Risk assessment studies on succinate dehydrogenase inhibitors, the new weapons in the battle to control Septoria leaf blotch in wheat. Mol Plant Pathol 2012, 13(3):263–275.

10. Hayes LE, Sackett KE, Anderson NP, Flowers MD, Mundt CC: Evidence of Selection for Fungicide Resistance in *Zymoseptoria tritici* Populations on Wheat in Western Oregon. Plant Dis 2016, 100(2):483–489.

11. Jess S, Kildea S, Moody A, Rennick G, Murchie AK, Cooke LR: European Union policy on pesticides: implications for agriculture in Ireland. Pest Manag Sci 2014, 70(11):1646–1654.

12. Wheat will not be a viable crop in Ireland by 2022, warn tillage experts

13. Kourelis J, van der Hoorn RAL: Defended to the Nines: 25 years of Resistance Gene Cloning Identifies Nine Mechanisms for R Protein Function. Plant Cell 2018.

14. Jones JDG, Dangl JL: The plant immune system. Nature 2006, 444(7117):323–329.

15. Swiderski MR, Innes RW: The Arabidopsis *PBS1* resistance gene encodes a member of a novel protein kinase subfamily. Plant J 2001, 26(1):101–112.

16. Sarris PF, Cevik V, Dagdas G, Jones JDG, Krasileva KV: Comparative analysis of plant immune receptor architectures uncovers host proteins likely targeted by pathogens. BMC Biol 2016, 14.

17. Cesari S, Bernoux M, Moncuquet P, Kroj T, Dodds PN: A novel conserved mechanism for plant NLR protein pairs: the “integrated decoy” hypothesis. Front Plant Sci 2014, 5.

18. Baggs E, Dagdas G, Krasileva KV: NLR diversity, helpers and integrated domains: making sense of the NLR IDentity. Curr Opin Plant Biol 2017, 38:59–67.

19. Noel L, Moores TL, van der Biezen EA, Parniske M, Daniels MJ, Parker JE, Jones JDG: Pronounced intraspecific haplotype divergence at the *RPP5* complex disease resistance locus of Arabidopsis. Plant Cell 1999, 11(11):2099–2111.

20. Seo E, Kim S, Yeom SI, Choi D: Genome-Wide Comparative Analyses Reveal the Dynamic Evolution of Nucleotide-Binding Leucine-Rich Repeat Gene Family among Solanaceae Plants. Front Plant Sci 2016, 7.

21. Kuang H, Woo SS, Meyers BC, Nevo E, Michelmore RW: Multiple genetic processes result in heterogeneous rates of evolution within the major cluster disease resistance genes in lettuce. Plant Cell 2004, 16(11):2870–2894.

22. Chavan S, Gray J, Smith SM: Diversity and evolution of *Rp1* rust resistance genes in four maize lines. Theor Appl Genet 2015, 128(5):985–998.

23. Cloutier S, McCallum BD, Loutre C, Banks TW, Wicker T, Feuillet C, Keller B, Jordan MC: Leaf rust resistance gene *Lr1*, isolated from bread wheat (*Triticum aestivum* L.) is a member of the large psr567 gene family. Plant Mol Biol 2007, 65(1-2):93–106.

24. Feuillet C, Travella S, Stein N, Albar L, Nublat A, Keller B: Map-based isolation of the leaf rust disease resistance gene *Lr10* from the hexaploid wheat (*Triticum aestivum* L.) genome. P Natl Acad Sci USA 2003, 100(25):15253–15258.

25. Huang L, Brooks SA, Li WL, Fellers JP, Trick HN, Gill BS: Map-based cloning of leaf rust resistance gene *Lr21* from the large and polyploid genome of bread wheat. Genetics 2003, 164(2):655–664.

26. Thind AK, Wicker T, Simkova H, Fossati D, Moullet O, Brabant C, Vrana J, Dolezel J, Krattinger SG: Rapid cloning of genes in hexaploid wheat using cultivar-specific long-range chromosome assembly. Nat Biotechnol 2017, 35(8):793–796.

27. Sanchez-Martin J, Steuernagel B, Ghosh S, Herren G, Hurni S, Adamski N, Vrana J, Kubalakova M, Krattinger SG, Wicker T et al: Rapid gene isolation in barley and wheat by mutant chromosome sequencing. Genome Biol 2016, 17(1):221.

28. Yahiaoui N, Srichumpa P, Dudler R, Keller B: Genome analysis at different ploidy levels allows cloning of the powdery mildew resistance gene *Pm3b* from hexaploid wheat. Plant J 2004, 37(4):528–538.

29. Hurni S, Brunner S, Buchmann G, Herren G, Jordan T, Krukowski P, Wicker T, Yahiaoui N, Mago R, Keller B: Rye *Pm8* and wheat *Pm3* are orthologous genes and show evolutionary conservation of resistance function against powdery mildew. Plant J 2013, 76(6):957–969.

30. SteuernagelB, Periyannan SK, Hernández-Pinzón I, Witek K, Rouse MN, Yu G, Hatta A, Ayliffe M, Bariana H, Jones JDG et al: Rapid cloning of disease-resistance genes in plants using mutagenesis and sequence capture. Nat Biotechnol 2016, in press.

31. Periyannan S, Moore J, Ayliffe M, Bansal U, Wang X, Huang L, Deal K, Luo M, Kong X, Bariana H et al: The gene *Sr33*, an ortholog of barley *Mla* genes, encodes resistance to wheat stem rust race Ug99. Science 2013, 341(6147):786–788.

32. Saintenac C, Zhang W, Salcedo A, Rouse MN, Trick HN, Akhunov E, Dubcovsky J: Identification of wheat gene *Sr35* that confers resistance to Ug99 stem rust race group. Science 2013, 341(6147):783–786.

33. Mago R, Zhang P, Vautrin S, Simkova H, Bansal U, Luo MC, Rouse M, Karaoglu H, Periyannan S, Kolmer J et al: The wheat *Sr50* gene reveals rich diversity at a cereal disease resistance locus. Nat Plants 2015, 1(12).

34. Marchal C, Zhang J, Zhang P, Fenwick P, Steuernagel B, Adamski NM, Boyd L, McIntosh R, Wulff BB, Berry S et al: BED-domain containing immune receptors confer diverse resistance spectra to yellow rust. bioRxiv 2018.

35. Consortium IWGS: Shifting the limits in wheat research and breeding using a fully annotated reference genome. under review 2018.

36. Bevan MW, Uauy C, Wulff BBH, Zhou J, Krasileva K, Clark MD: Genomic innovation for crop improvement. Nature 2017, 543(7645):346–354.

37. Brenchley R, Spannagl M, Pfeifer M, Barker GLA, D’Amore R, Allen AM, McKenzie N, Kramer M, Kerhornou A, Bolser D et al: Analysis of the bread wheat genome using whole-genome shotgun sequencing. Nature 2012, 491(7426):705–710.

38. International Wheat Genome Sequencing C: A chromosome-based draft sequence of the hexaploid bread wheat (*Triticum aestivum*) genome. Science 2014, 345(6194):1251788.

39. Chapman JA, Mascher M, Buluc A, Barry K, Georganas E, Session A, Strnadova V, Jenkins J, Sehgal S, Oliker L et al: A whole-genome shotgun approach for assembling and anchoring the hexaploid bread wheat genome. Genome Biology 2015, 16.

40. Clavijo BJ, Venturini L, Schudoma C, Accinelli GG, Kaithakottil G, Wright J, Borrill P, Kettleborough G, Heavens D, Chapman H et al: An improved assembly and annotation of the allohexaploid wheat genome identifies complete families of agronomic genes and provides genomic evidence for chromosomal translocations. Genome Res 2017, 27(5):885–896.

41. Zimin AV, Puiu D, Hall R, Kingan S, Clavijo BJ, Salzberg SL: The first near-complete assembly of the hexaploid bread wheat genome, *Triticum aestivum*. Gigascience 2017, 6(11).

42. Ramírez-González RH, Borill P, Lang D, Harrington SA, Brinton J, Venturini L, Davey M, Jacobs J, van Ex F, Pasha A et al: The transcriptional landscape of polyploid wheat. under review 2018.

43. Luo MC, Gu YQ, Puiu D, Wang H, Twardziok SO, Deal KR, Huo NX, Zhu TT, Wang L, Wang Y et al: Genome sequence of the progenitor of the wheat D genome *Aegilops tauschii*. Nature 2017, 551(7681):498-+.

44. Zhao GY, Zou C, Li K, Wang K, Li TB, Gao LF, Zhang XX, Wang HJ, Yang ZJ, Liu X et al: The *Aegilops tauschii* genome reveals multiple impacts of transposons. Nat Plants 2017, 3(12):946–955.

45. Avni R, Nave M, Barad O, Baruch K, Twardziok SO, Gundlach H, Hale I, Mascher M, Spannagl M, Wiebe K et al: Wild emmer genome architecture and diversity elucidate wheat evolution and domestication. Science 2017, 357(6346):93–96.

46. Andolfo G, Jupe F, Witek K, Etherington GJ, Ercolano MR, Jones JDG: Defining the full tomato NB-LRR resistance gene repertoire using genomic and cDNA RenSeq. BMC Plant Biol 2014, 14.

47. Steuernagel B, Jupe F, Witek K, Jones JDG, Wulff BBH: NLR-parser: rapid annotation of plant NLR complements. Bioinformatics 2015, 31(10):1665–1667.

48. Jupe F, Pritchard L, Etherington GJ, MacKenzie K, Cock PJA, Wright F, Sharma SK, Bolser D, Bryan GJ, Jones JDG et al: Identification and localisation of the NB-LRR gene family within the potato genome. BMC Genomics 2012, 13.

49. Denoeud F, Carretero-Paulet L, Dereeper A, Droc G, Guyot R, Pietrella M, Zheng CF, Alberti A, Anthony F, Aprea G et al: The coffee genome provides insight into the convergent evolution of caffeine biosynthesis. Science 2014, 345(6201):1181–1184.

50. Jiao YP, Peluso P, Shi JH, Liang T, Stitzer MC, Wang B, Campbell MS, Stein JC, Wei XH, Chin CS et al: Improved maize reference genome with single-molecule technologies. Nature 2017, 546(7659):524-+.

51. Ming R, Hou SB, Feng Y, Yu QY, Dionne-Laporte A, Saw JH, Senin P, Wang W, Ly BV, Lewis KLT et al: The draft genome of the transgenic tropical fruit tree papaya (*Carica papaya* Linnaeus). Nature 2008, 452(7190):991–U997.

52. Schmutz J, Cannon SB, Schlueter J, Ma JX, Mitros T, Nelson W, Hyten DL, Song QJ, Thelen JJ, Cheng JL et al: Genome sequence of the palaeopolyploid soybean (vol 463, pg 178, 2010). Nature 2010, 465(7294):120–120.

53. Xu X, Pan SK, Cheng SF, Zhang B, Mu DS, Ni PX, Zhang GY, Yang S, Li RQ, Wang J et al: Genome sequence and analysis of the tuber crop potato. Nature 2011, 475(7355):189–U194.

54. Sato S, Tabata S, Hirakawa H, Asamizu E, Shirasawa K, Isobe S, Kaneko T, Nakamura Y, Shibata D, Aoki K et al: The tomato genome sequence provides insights into fleshy fruit evolution. Nature 2012, 485(7400):635–641.

55. Vogel JP, Garvin DF, Mockler TC, Schmutz J, Rokhsar D, Bevan MW, Barry K, Lucas S, Harmon-Smith M, Lail K et al: Genome sequencing and analysis of the model grass *Brachypodium distachyon*. Nature 2010, 463(7282):763–768.

56. Loutre C, Wicker T, Travella S, Galli P, Scofield S, Fahima T, Feuillet C, Keller B: Two different CC-NBS-LRR genes are required for *Lr10*-mediated leaf rust resistance in tetraploid and hexaploid wheat. Plant J 2009, 60(6):1043–1054.

57. Parlange F, Roffler S, Menardo F, Ben-David R, Bourras S, McNally KE, Oberhaensli S, Stirnweis D, Buchmann G, Wicker T et al: Genetic and molecular characterization of a locus involved in avirulence of *Blumeria graminis* f. sp *tritici* on wheat *Pm3* resistance alleles. Fungal Genet Biol 2015, 82:181–192.

58. Isidore E, Scherrer B, Chalhoub B, Feuillet C, Keller B: Ancient haplotypes resulting from extensive molecular rearrangements in the wheat A genome have been maintained in species of three different ploidy levels. Genome Res 2005, 15(4):526–536.

59. Meyers BC, Dickerman AW, Michelmore RW, Sivaramakrishnan S, Sobral BW, Young ND: Plant disease resistance genes encode members of an ancient and diverse protein family within the nucleotide-binding superfamily. Plant J 1999, 20(3):317–332.

60. Seeholzer S, Tsuchimatsu T, Jordan T, Bieri S, Pajonk S, Yang WX, Jahoor A, Shimizu KK, Keller B, Schulze-Lefert P: Diversity at the *Mla* Powdery Mildew Resistance Locus from Cultivated Barley Reveals Sites of Positive Selection. Mol Plant-Microbe Interact 2010, 23(4):497–509.

61. Halterman D, Zhou FS, Wei FS, Wise RP, Schulze-Lefert P: The MLA6 coiled-coil, NBS-LRR protein confers *AvrMla6*-dependent resistance specificity to *Blumeria graminis* f. sp *hordei* in barley and wheat. Plant J 2001, 25(3):335–348.

62. Wei FS, Gobelman-Werner K, Morroll SM, Kurth J, Mao L, Wing R, Leister D, Schulze-Lefert P, Wise RP: The *Mla* (powdery mildew) resistance cluster is associated with three NBS-LRR gene families and suppressed recombination within a 240-kb DNA interval on chromosome 5S (1HS) of barley (vol 153, pg 1929, 1999). Genetics 2000, 154(2):953–953.

63. Bailey PC, Schudoma C, Jackson W, Baggs E, Dagdas G, Haerty W, Moscou M, Krasileva KV: Dominant integration locus drives continuous diversification of plant immune receptors with exogenous domain fusions. Genome Biol 2018, 19(1):23.

64. Stein JC, Yu Y, Copetti D, Zwickl DJ, Zhang L, Zhang CJ, Chougule K, Gao DY, Iwata A, Goicoechea JL et al: Genomes of 13 domesticated and wild rice relatives highlight genetic conservation, turnover and innovation across the genus *Oryza*. Nat Genet 2018, 50(2):285-+.

65. Narusaka M, Shirasu K, Noutoshi Y, Kubo Y, Shiraishi T, Iwabuchi M, Narusaka Y: *RRS1* and *RPS4* provide a dual Resistance-gene system against fungal and bacterial pathogens. Plant J 2009, 60(2):218–226.

66. Jupe F, Witek K, Verweij W, Sliwka J, Pritchard L, Etherington GJ, Maclean D, Cock PJ, Leggett RM, Bryan GJ et al: Resistance gene enrichment sequencing (RenSeq) enables reannotation of the *NB-LRR* gene family from sequenced plant genomes and rapid mapping of resistance loci in segregating populations. Plant J 2013, 76(3):530–544.

67. Yi HK, Richards EJ: Gene Duplication and Hypermutation of the Pathogen Resistance Gene *SNC1* in the Arabidopsis bal Variant. Genetics 2009, 183(4):1227–1234.

68. Choulet F, Alberti A, Theil S, Glover N, Barbe V, Daron J, Pingault L, Sourdille P, Couloux A, Paux E et al: Structural and functional partitioning of bread wheat chromosome 3B. Science 2014, 345(6194).

69. Dangl JL, Horvath DM, Staskawicz BJ: Pivoting the Plant Immune System from Dissection to Deployment. Science 2013, 341(6147):746–751.

70. Kombrink A, Sanchez-Vallet A, Thomma B: The role of chitin detection in plant-pathogen interactions. Microb Infect 2011, 13(14-15):1168–1176.

71. Takai R, Isogai A, Takayama S, Che FS: Analysis of Flagellin Perception Mediated by flg22 Receptor OsFLS2 in Rice. Mol Plant-Microbe Interact 2008, 21(12):1635–1642.

72. Schoonbeek HJ, Wang HH, Stefanato FL, Craze M, Bowden S, Wallington E, Zipfel C, Ridout CJ: Arabidopsis EF-Tu receptor enhances bacterial disease resistance in transgenic wheat. New Phytol 2015, 206(2):606–613.

73. Bailey TL, Gribskov M: Combining evidence using p-values: application to sequence homology searches. Bioinformatics 1998, 14(1):48–54.

74. Letunic I, Doerks T, Bork P: SMART: recent updates, new developments and status in 2015. Nucleic Acids Res 2015, 43(D1):D257–D260.

75. Zhang Z, Schwartz S, Wagner L, Miller W: A greedy algorithm for aligning DNA sequences. J Comput Biol 2000, 7(1-2):203–214.

76. Sonnhammer ELL, Durbin R: A dot-matrix program with dynamic threshold control suited for genomic DNA and protein sequence analysis (Reprinted from Gene Combis, vol 167, pg GC1-GC10, 1996). Gene 1995, 167(1-2):Gc1–Gc10.

77. Lee E, Helt GA, Reese JT, Munoz-Torres MC, Childers CP, Buels RM, Stein L, Holmes IH, Elsik CG, Lewis SE: Web Apollo: a web-based genomic annotation editing platform. Genome Biology 2013, 14(8).

78. Larkin MA, Blackshields G, Brown NP, Chenna R, McGettigan PA, McWilliam H, Valentin F, Wallace IM, Wilm A, Lopez R et al: Clustal W and clustal X version 2.0. Bioinformatics 2007, 23(21):2947–2948.

79. Price MN, Dehal PS, Arkin AP: FastTree 2-Approximately Maximum-Likelihood Trees for Large Alignments. PloS one 2010, 5(3).

80. Letunic I, Bork P: Interactive tree of life (iTOL) v3: an online tool for the display and annotation of phylogenetic and other trees. Nucleic Acids Res 2016, 44(W1):W242–W245.

81. Eddy SR: Accelerated Profile HMM Searches. PLoS Comp Biol 2011, 7(10).

82. Bray NL, Pimentel H, Melsted P, Pachter L: Near-optimal probabilistic RNA-seq quantification. Nat Biotechnol 2016, 34(5):525–527.

